# Mapping the temporal and spatial dynamics of the human endometrium *in vivo* and *in vitro*

**DOI:** 10.1101/2021.01.02.425073

**Authors:** Luz Garcia-Alonso, Louis-François Handfield, Kenny Roberts, Konstantina Nikolakopoulou, Ridma C. Fernando, Lucy Gardner, Ben Woodhams, Anna Arutyunyan, Krzysztof Polanski, Regina Hoo, Carmen Sancho-Serra, Tong Li, Kwasi Kwakwa, Elizabeth Tuck, Vitalii Kleshchevnikov, Aleksandra Tarkowska, Tarryn Porter, Cecilia Icoresi Mazzeo, Stijn van Dongen, Monika Dabrowska, Vasyl Vaskivskyi, Krishna T. Mahbubani, Jong-eun Park, Mercedes Jimenez-Linan, Lia Campos, Vladimir Kiselev, Cecilia Lindskog, Paul Ayuk, Elena Prigmore, Michael R Stratton, Kourosh Saeb-Parsy, Ashley Moffett, Luiza Moore, Omer A. Bayraktar, Sarah A. Teichmann, Margherita Y. Turco, Roser Vento-Tormo

## Abstract

The endometrium, the mucosal lining of the uterus, undergoes dynamic changes throughout the menstrual cycle in response to ovarian hormones. We have generated single-cell and spatial reference maps of the human uterus and 3D endometrial organoid cultures. We dissect the signalling pathways that determine cell fate of the epithelial lineages in the lumenal and glandular microenvironments. Our benchmark of the endometrial organoids highlights common pathways regulating the differentiation of secretory and ciliated lineage *in vivo* and *in vitro*. We show *in vitro* that downregulation of WNT or NOTCH pathways increases the differentiation efficiency along the secretory and ciliated lineages, respectively. These mechanistic insights provide a platform for future development of treatments for a range of common endometrial disorders including endometriosis and carcinoma.

---

The human endometrium, the mucosal lining of the uterus, is the site of implantation that provides nutritional support to the placenta throughout pregnancy. Unlike other mucosal tissues it undergoes dynamic, cyclical changes of shedding, regeneration and differentiation throughout reproductive life coordinated by the hypothalamic-pituitary-ovarian axis. Endometrial dysfunction underpins many common disorders including abnormal uterine bleeding, infertility, miscarriage, pre-eclampsia, endometriosis and endometrial carcinoma that collectively affect many women across the world (*1*–*5*). Throughout reproductive years the functional upper layer of the endometrium, the *stratum functionalis*, is shed at menstruation. The subsequent tissue repair and proliferation are driven by the rising levels of estrogen, secreted by the ovarian follicle during the first half of the menstrual cycle (proliferative phase). Following ovulation, progesterone, produced by the *corpus luteum*, induces the secretory phase during which the initial changes of decidualisation occur. Menstruation and spontaneous decidualisation are unique to higher simian primates (*6*–*8*). Thus, dissecting the mechanisms that regulate cellular differentiation across the menstrual cycle in humans is crucial for understanding how normal endometrium is regulated.

Essential to endometrial function are the lumenal and glandular epithelia, composed of a mixture of ciliated and secretory cells. The lumenal epithelium is the site of embryo attachment. This epithelium covers the endometrial surface and, from it, invaginations into the stroma form the glands. Glandular secretions are rich in growth factors and lipids necessary for placental growth (*9*). We have established a three-dimensional (3D) *in vitro* organoid culture model of human endometrial epithelium (*10*, *11*). These organoids are generated from dissociated endometrial tissue, they retain the morphology, function and gene signature of the tissue *in vivo* and respond functionally to ovarian hormones with differentiation into ciliated and secretory cells. However, in our initial study, we did not perform a systematic, quantitative comparison of endometrial organoids at a single cell level with epithelial cell states *in vivo*. This is needed to confirm their suitability for exploring the cell pathways and processes involved in normal and pathological endometrial function.

The explosion in spatial transcriptomics technologies (*12*–*15*) provides a unique opportunity to resolve tissue architecture in conjunction with the underlying cellular interactions. The spatial arrangement of cells is key to understanding a morphologically complex tissue such as the endometrium, where a cell’s function may differ depending on signals it receives from neighbouring cells (*16*). Many spatially-resolved transcriptomics methods are not quite at single-cell resolution, and rely on the computational integration of coupled single-cell (or single nuclei) transcriptomes to achieve this level of detail (*17*–*19*). These genomic technologies are the basis of the Human Cell Atlas initiative, which aims to map all cells in the human body (*20*).

In this study, by using single-cell and spatial transcriptional profiling, we interrogate the cellular states and spatial localisation of human endometrial cells during the proliferative and secretory phases of the menstrual cycle in women of reproductive age. We develop CellPhoneDB v3.0 to measure intercellular communication taking into account spatial coordinates of cells, and use this tool to define cell signalling in both lumenal and glandular epithelial microenvironments. We define a complementary role for WNT and NOTCH signalling in regulating differentiation towards the two main epithelial lineages (ciliated and secretory). We profile 3D endometrial organoids at single-cell resolution to characterise their hormonal responses *in vitro* and design a novel computational toolkit to compare the results with those observed *in vivo* in order to benchmark this model system. Finally, by modulating WNT and NOTCH pathways in the organoid cultures, we develop lineage-specific endometrial epithelial cells, and define the molecular events involved in their response to ovarian hormones.

## Results

### A single-cell map of the full thickness human uterus

To generate a cellular map of the human endometrium that accounts for the temporal and spatial changes across the menstrual cycle, uterine samples were analysed by single-cell transcriptomics (single-cell RNA sequencing (scRNA-seq) and single-nuclei RNA sequencing (snRNA-seq)) alongside spatial transcriptomics methods (10x Genomics Visium slides and high-resolution microscopy) (**Fig. 1A, Fig. S1A**). Two different types of samples from women of reproductive age were integrated in our analysis: superficial endometrial biopsies from live donors (n=3) and full-thickness uterine tissue from donors who died of non-gynaecological causes (n=8) (**Table S1**). The latter approach allows exploration of cell signatures in the basal layer of the endometrium and the myometrium for the first time.

**Fig. 1.**
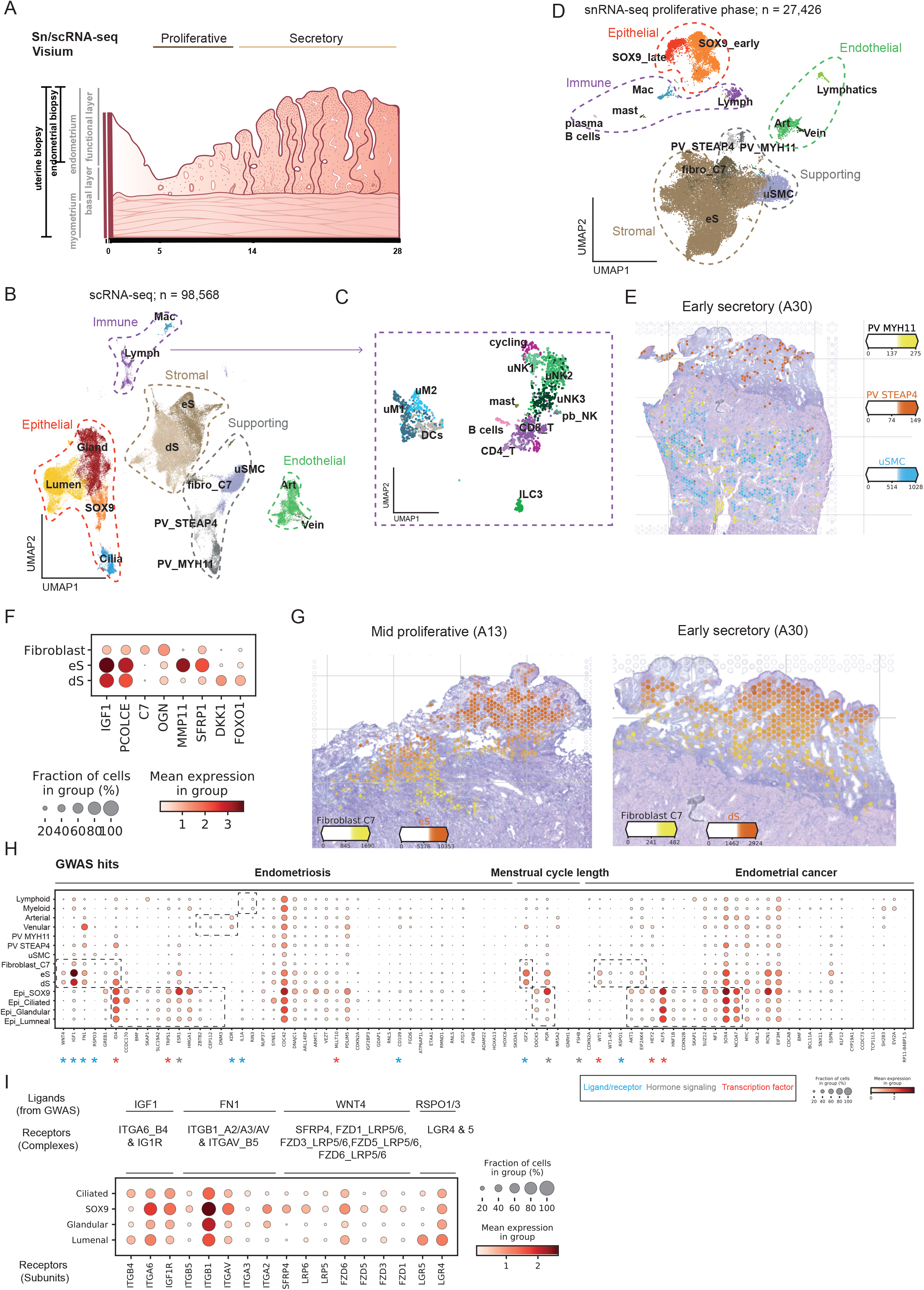
Single-cell profiling of the human uterus. **(A)** Schematic illustration of the human uterus showing the different layers and their morphological changes across the phases of the menstrual cycle. **(B)** UMAP (uniform manifold approximation and projection) projections of scRNA-seq data from a total of 15 individuals. **(C)** UMAP of sub-clustered immune populations. **(D)** UMAP projections of snRNA-seq data from a total of 4 individuals. (**E)** Estimated amount of mRNA (colour intensity) contributed by each cell population to each spot (colour) shown over the H&E image of the secretory endometrium (sample 152811 -A30-). (**F)** Dot plot showing log_2_-transformed expression of genes expressed in the fibroblast and stromal subsets. **(G)** Estimated amount of mRNA (colour intensity) contributed by each cell population to each spot (colour) shown over the H&E image of the proliferative (sample 152810 -A13-) and secretory (sample 152811 -A30-) endometrium. **(H)** Dot plot showing log_2_-transformed expression of genes associated with endometriosis and regulation of the length of the menstrual cycle by GWAS, projected onto all uterine cell states. **(I)** Dot plot showing log_2_-transformed expression of receptor subunits that can potentially bind the ligands linked to endometriosis and endometrial cancer GWAS loci. Lumen = lumenal; Gland = glandular; uSMC = uterine smooth muscle cell; PV = perivascular; eS = non-decidualised endometrial stromal cells; dS = decidualised endometrial stromal cells; art = artery; fibro = fibroblasts; uM = uterine macrophages; uNK = uterine Natural Killer cells, T = T cells, ILC = Innate lymphoid cells, DC = Dendritic cells; scRNA-seq = single-cell RNA sequencing; snRNA-seq = single-nuclei RNA sequencing.

We were able to generate a uterine map of 98,568 cells from fifteen individuals by integrating our dataset with previous scRNA-seq data obtained from superficial endometrial biopsies (*21*) (**Fig. 1B, Fig. S1B-D, Fig. S2A-E, Table S2**). We identify fourteen clusters that were assigned cell identity based on their expression of known markers (**Fig. S2F**). These clusters can be grouped into five main cellular categories: (i) immune (lymphoid and myeloid), (ii) epithelial (*SOX9+*, lumenal, glandular and ciliated), (iii) endothelial (arterial and venular) (iv) supporting (perivascular (PV), smooth muscle cells (SMC) and fibroblasts (C7)), and (v) stromal (non-decidualised endometrial (eS) and decidualised endometrial (dS)). *SOX9+* epithelial cells and eS are characteristic of the regenerating proliferative phase (**Fig. S2D**). Sub-clustering of immune cells allows us to resolve their heterogeneity, including identification of the three uterine Natural Killer (NK) cell subsets we have previously defined in the early pregnant uterus (*22*) (**Fig. 1C, Fig. S2G**). SnRNA-seq data from four additional full-thickness uterine samples of proliferative endometrium confirms the populations found in the poorly-studied regenerative stage (**Fig. 1D**, **Fig. S3A-E**). We also found lymphatic endothelial cells enriched in these samples; these are likely to be present in the myometrium (**Fig. S3D**).

To systematically map the location of the cell types identified by scRNA-seq within the endometrium and myometrium, we used Visium Spatial Transcriptomics technology. We examined four full-thickness uterine samples in the proliferative and secretory phases from two individuals (**Fig. S4A-E**). After integrating single-cell transcriptomics and Visium data using our recently developed cell2location algorithm (*17*), cell states were mapped to the endometrium and/or the myometrium. We identify specific PV cells: PV-MYH11 are characteristic of myometrium whilst PV-STEAP4 are only present in the endometrium (**Fig. 1E, Fig. S4F**). In addition, we find that a novel population of fibroblasts identified by scRNA-seq (fibroblasts C7) is enriched in the basal layer of the endometrium in both proliferative and secretory phases (**Fig. 1F-G, Fig. S4G**).

Altogether, our analysis yields a comprehensive catalogue of the major subsets of uterine cells together with their cellular position in endometrium and myometrium. We have made an open-source web server available at www.reproductivecellatlas.org.

### Epithelial cells are the main target of endometrial disorders

Disorders in endometrial function have a profound impact on women’s health and reproductive outcomes. There has been little progress in the study of endometrial disorders over the past decade, partly due to the challenges in analysing this highly dynamic and complex tissue. In order to identify the cells involved in these pathologies, we interrogated the genetic variants associated with endometriosis (*23*), endometrial cancer (*24*, *25*) and menstrual cycle length (*26*) in our scRNA-seq data of the human uterus. To link the genetic associations found with molecular mechanisms, we classified the GWAS hits into ligand-receptor interactions, hormonal signalling pathways and transcription factors (TFs) (**Fig. 1H**).

We find that the majority of the GWAS hits in endometriosis and endometrial cancer are associated with genes expressed by epithelial cells. A significant number of these genes are characteristic of the proliferative phase (*SOX9* subset), including those involved in the regulation of the cell cycle and apoptosis. These are: cell cycle regulatory genes, *AKT1* (endometrial cancer); TFs, *TRPS1* (endometriosis) and *KLF5* (endometrial cancer); and hormonal receptors, *ESR1* (endometriosis) and *PGR* (regulation of menstrual cycle length).

Many of these GWAS loci encode for ligands and receptors highlighting the importance of cell-cell communication within the endometrium. The majority of the ligands are expressed by stromal cells and fibroblast C7, with their cognate receptors differentially expressed by epithelial subsets (**Fig. 1I**). *WNT4* (endometriosis), expressed by stromal cells, can potentially bind WNT receptors highly expressed by epithelial cells in the proliferative stage. R-spondins, which activate the WNT pathway after binding Lgr4/5/6 receptors, are associated with endometrial cancer (*RSPO1*) and endometriosis *(RSPO3*) and are expressed by fibroblast C7. Thus, our analysis suggests that a major driver of endometrial disease is a dysfunctional dialogue between epithelial and stromal cells.

### Temporal and spatial analysis of the endometrial epithelium defines the cellular landscape of the proliferative phase

We therefore focussed next on the two main lineages of endometrial epithelial cells, secretory and ciliated, across the menstrual cycle and analysed these subsets individually (**Fig. 2A**). Epithelial cells are classified into four main groups based on their marker expression: (i) *SOX9* populations, enriched in the proliferative phase and expressing genes characteristic of rising estrogen levels (*MMP7, ESR1*), ciliated cells (*PIFO*, *TPPP3*), (iii) lumenal cells (*LGR5*) and (iv) glandular cells (*SCGB2A2*) (**Fig. 2B-D, Fig. S5A-B**). A fraction of the glandular epithelial cells express molecules characteristic of uterine milk in the secretory stage *(PAEP, CXCL8).*

**Fig. 2.**
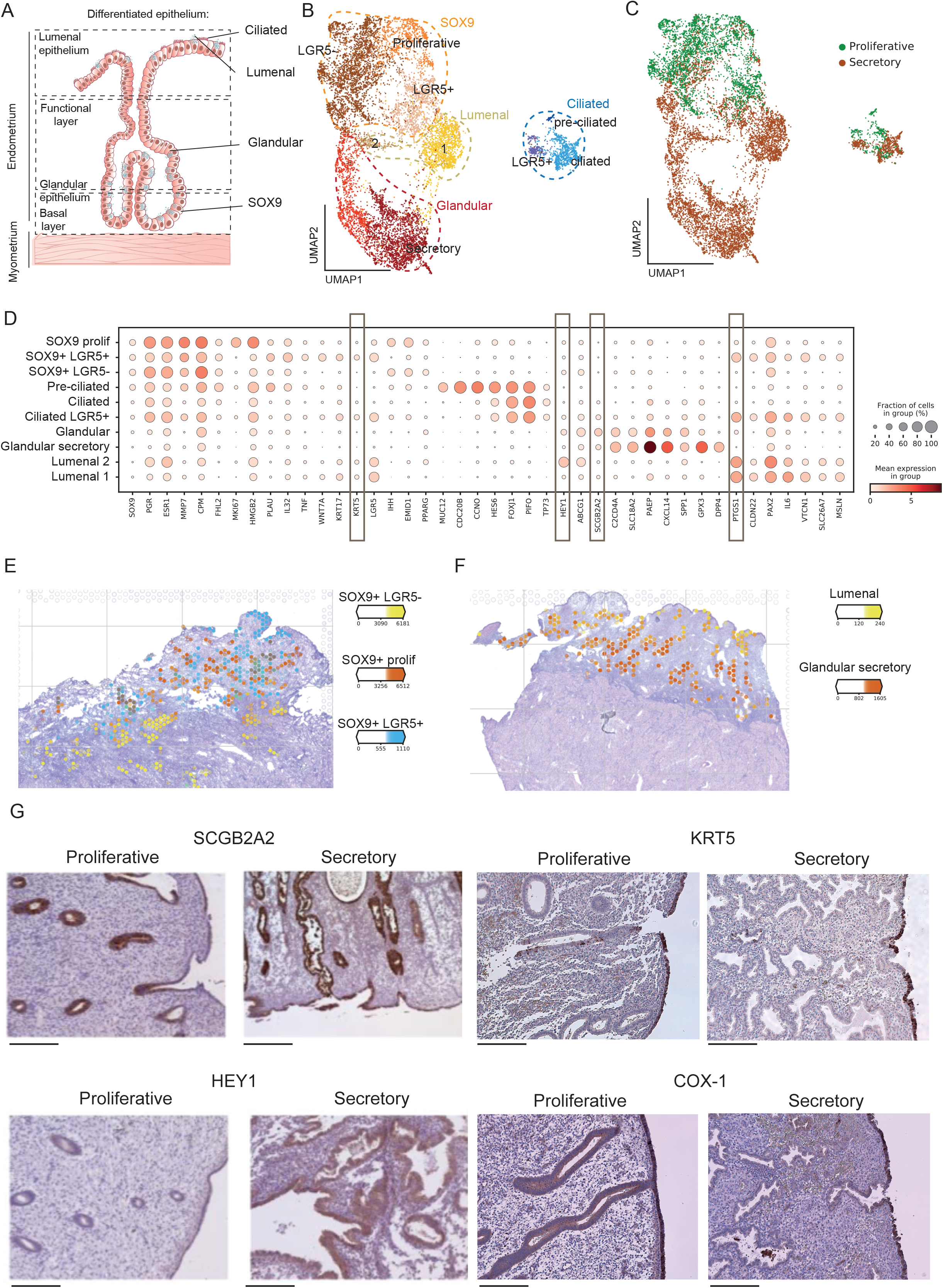
Temporal and spatial dynamics of endometrial epithelial cells. **(A)** Schematic illustration of epithelial subsets in the endometrium highlighting the anatomical location of the glandular and lumenal epithelia. **(B)** UMAP of sub-clustered and sub-sampled epithelial populations. **(C)** UMAP of sub-sampled epithelial populations coloured by their menstrual stage. **(D)** Dot plot showing the log_2_-transformed expression of genes characteristic of each epithelial subsets. **(E)** Number of mRNA molecules per spot (colour intensity) confidently assigned to each epithelial subpopulation (colour) in the proliferative phase (A13, 152810 slide). **(F)** Number of mRNA molecules per spot (colour intensity) confidently assigned to each epithelial subpopulation (colour) in the early-proliferative phase (A30, 152807 slide). **(G)** Validation of KRT5, COX-1 (marker of lumenal populations), SCGB2A2 (marker of glandular population), HEY1 (marker of secretory cells) with immunohistochemistry (IHC) in endometrial tissue (proliferative and secretory phases). Nuclei are counterstained with hematoxylin. Scale bars = 250 μm. Representative images of four proliferative and secretory endometrial samples from eight different donors.

The composition of proliferative phase epithelial cells has been largely unexplored as samples are less frequently taken during this time. Our integrative scRNA-seq maps resolve three clusters within the *SOX9* population: (i) *SOX9+LGR5+* cells, which express markers such as *IL32* or *TNF*, (ii) *SOX9+LGR5−*, expressing *PPARG*, (iii) proliferative *SOX9+* which includes both *LGR5+* and *LGR5−* cycling cells (*i.e.*, cells in G2M/S phase). By integrating scRNA-seq and Visium data (**Fig. 2E, Fig. S5C**), we define specific spatial coordinates for each subset: (i) non-cycling *SOX9+LGR5+* cells are enriched in the surface epithelium (ii) non-cycling *SOX9+LGR5−* cells are located in the basal glands (iii) cycling *SOX9+* cells map to glands in the regenerating superficial layer. Single-molecule fluorescence in situ hybridization (smFISH) with RNAscope probes locates the *SOX9+LGR5+* markers (*LGR5, WNT7A*) to the surface epithelium during the proliferative phase (**Fig. S6A-B**).

Ciliated cells are present in both the proliferative and secretory phase, but, as expected, *PAEP* secretory cells are only present following ovulation (**Fig. 2B-C, Fig. S6C)**. This indicates that estrogen alone can induce ciliary differentiation while secretory differentiation depends on the addition of progesterone. During the proliferative phase, in addition to *FOXJ1, PIFO, TP73* ciliated cells, we define a novel subset of pre-ciliated cells that express cell cycle genes (*CDC20B, CCNO*), *MUC12* and *HES6* (**Fig. 2D)**. We also find a subset of lumenal epithelial cells that co-express FOXJ1 (ciliated cells) and *LGR5* (lumenal cells) using smFISH (**Fig. S6D)**. This finding confirms the location of *FOXJ1, LGR5* cells found in our scRNA-seq data.

After ovulation, secretion of progesterone induces the differentiation of *SOX9+* cells into specialised secretory cells that can be found at the surface and in glands. Specific transcriptomics signatures and markers that can distinguish between lumenal and glandular differentiated epithelial subsets have been lacking. The identification of these subsets in our scRNA-seq data, confirmed by integration with spatial transcriptomics data (**Fig. 2F, Fig. S5D-E)**, now reveals novel markers that are validated at the protein level (**Fig. 2G**). There is enriched expression of both COX1 (encoded by *PTGS1*) and KRT5 in the lumenal epithelium and SCGB2A2 in the glandular epithelium. The NOTCH target HEY1 is induced in the glands and lumen after ovulation.

### Microenvironments modulate epithelial cell identity

We next investigated the TFs that regulate epithelial changes throughout the menstrual cycle by exploring their expression and that of their consensus target genes (*27*) (**Fig. 3A, Table S3**). Our analysis reveals high activity of WNT targets (e.g. *FOXJ1*) in the ciliated epithelium. In contrast, glandular subsets show high expression for TF induced by WNT inhibition (e.g. *CSRNP1* and *FOXO1*) and NOTCH activation (e.g. *HES1* and *HEY1*). This suggests different roles for NOTCH and WNT in shaping identity and function of ciliated and secretory cells.

**Fig. 3.**
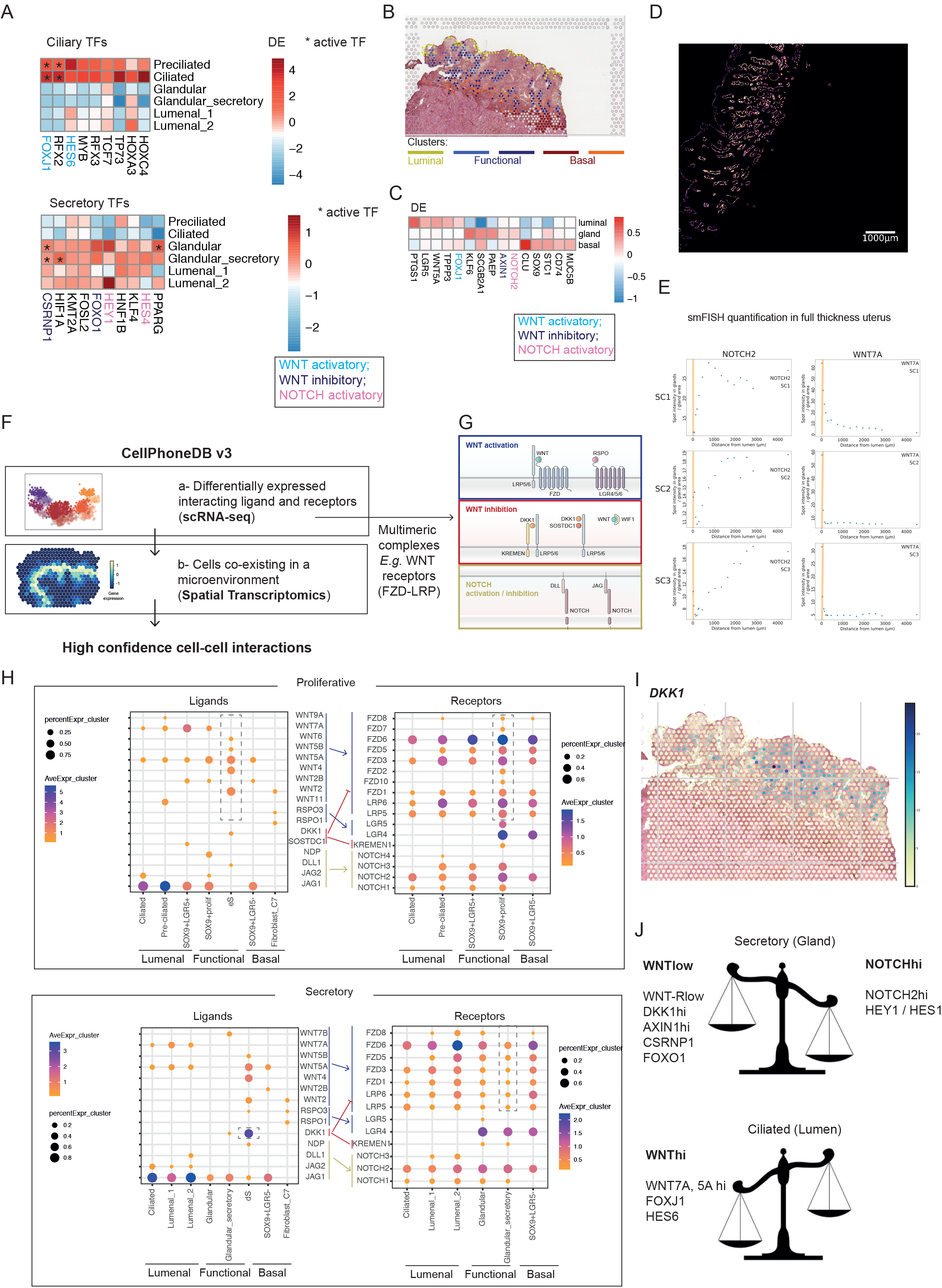
Cell signalling in the glands and lumen. **(A)** Heatmaps showing TFs differentially expressed (DE) in Ciliated (top) and Secretory (bottom) epithelial lineages. Color proportional to log-fold change, with asterisk (*) highlighting TFs whose targets are also differentially expressed (*i.e.* differentially activated TFs). **(B)** Unbiased clustering of epithelial subsets using Visium data. Spot colours represent cluster assignment based on Louvain clustering of spots assigned to epithelial subsets. Spots assigned to one of the clusters (represented in light grey in the figure) were excluded from the analysis due to the low percentage of epithelial cells in the spot after visual inspection. **(C)** Heatmap showing log-fold change of DE genes between the three main clusters defining the lumenal, functional and basal epithelial regions. **(D)** High-resolution large-area imaging of a representative uterine section of the secretory phase, with intensity proportional to smFISH signal for *NOTCH2*. Scale bar = 1000 μm. **(E)** smFISH quantification in three full thickness secretory phase uterine samples. SC = Secretory Section of the uterus. Plots represent smFISH spot intensity in glands divided by gland area at increasing distances from the lumen. Approximate lumen range is marked in yellow. **(F)** Adaptation of our cell-cell communication pipeline to consider spatial dynamics of cells. The tool is available at https://github.com/Ventolab/Cellphonedbv3. **(G)** Schematic illustration of receptors and ligands involved in WNT and NOTCH signalling. **(H)** Dot plots showing expression of ligands in the epithelial, stromal and fibroblast populations and cognate receptors in the epithelial subsets. Only significant interactions (FC >0.02 and DFR <0.005 are represented). Colour of the arrows correspond to the pathways those ligand/receptor partners are involved as shown in **(G)**. (**I)** Estimated proportions of *DKK1* coming from dS in the early-proliferative phase (A30, 152807 slide). (**J)** Schematic illustration of our model for NOTCH and WNT regulation in the glandular and lumenal epithelium. eS = non-decidualised endometrial stromal cell; dS = decidualised endometrial stromal cell.

To investigate the cell signals operating in the lumenal and glandular microenvironments that could influence differentiation into ciliated and secretory lineages, we used spatial transcriptomics and performed clustering on the 10x Genomics Visium spots assigned to epithelial subsets. We resolve five clusters corresponding to cells in the lumenal (one cluster), functional (two clusters) and basal (two clusters) layers (**Fig. 3B**). In addition to cell-type specific markers, signatures of WNT and NOTCH signalling pathways are present in distinct endometrial regions (**Fig. 3C, Table S4**). Genes involved in the WNT pathway, *FOXJ1*, *WNT5A*, and *LGR5*, are highly expressed at the lumenal surface whilst *NOTCH2* is enriched in glands in the functional layer. To validate expression of *NOTCH2* and *WNT7A* in these compartments, we stained uterine tissue with smFISH probes for both genes alongside EPCAM using immunohistochemistry (IHC) (**Fig. 3D**). Lumenal and glandular epithelial cells were classified automatically based on EPCAM expression and the distance of the signal away from the lumen was then measured (**see Methods**). Our results show *NOTCH2* expression increases in glands moving away from the lumen while *WNT7A* expression is higher in the lumenal epithelium compared to glands (**Fig. 3E**).

To investigate how surrounding cells may shape signalling in the surface and glandular compartments, we developed CellPhoneDBv3, a new version of our cell-cell communication pipeline that takes into account spatial cellular coordinates when mapping ligand-receptor pairs (*28*) (**see Methods**) (**Fig. 3F**). CellPhoneDB considers the multimeric composition of the majority of ligands and receptors, obviously relevant for the complex regulation of WNT signalling (**Fig. 3G**). We define four endometrial microenvironments based on the cellular coordinates provided by cell2location: i) lumenal - pre-ciliated, ciliated and *SOX9+LGR5+* epithelium (proliferative phase) and ciliated and lumenal (secretory phase), ii) functional - *SOX9+* proliferative epithelium and eS (proliferative phase) and glandular and dS (secretory phase), iii) vascular - PV-STEAP4 and endothelial vessels, iv) basal - *SOX9+LGR5−* and basal fibroblasts. We ran CellPhoneDB on each of the microenvironments (**Table S5**) and found significant epithelial-stromal interactions (0.02 > FC / 0.005 < FDR).

The NOTCH ligand *JAG1* is strongly expressed by all epithelial subsets. In contrast, WNT ligands are expressed by both epithelial and stromal cells (**Fig. 3H**); the latter express WNT agonists that can potentially bind the cognate WNT receptors expressed by all epithelial subsets during the proliferative phase (**Fig. 3H**). Focussing on genes that are differentially expressed following ovulation, glandular secretory subsets show a dramatic decline in WNT receptor expression, potentially limiting the activity of this pathway (**Fig. 3H**). In addition, decidualised stromal cells express significantly higher levels of *DKK1,* a potent inhibitor of the WNT pathway, than their non-decidualised counterparts. Expression of *DKK1* surrounding the glands of the secretory endometrium is also found in spatial transcriptomics (**Fig. 3I**). Overall, these findings strongly suggest that WNT signalling is inhibited in the secretory cell lineages, meaning that NOTCH signalling will then dominate (**Fig. 3J**).

### Endometrial organoids recapitulate transcriptional programs induced by hormonal stimulation

To test our predictions on the potential roles of WNT and NOTCH signalling pathways on endometrial epithelium *in vitro,* we first profiled endometrial organoids at a single cell level to benchmark this model system against our *in vivo* data. We derived organoids from three different donors and primed them as previously described with estrogen (E2) for 48 hours, followed by stimulation with progesterone (P4), prolactin (PRL) and cAMP in the presence of E2 for four days (*10*) (**Fig. 4A, Table S6-7**).

**Fig. 4.**
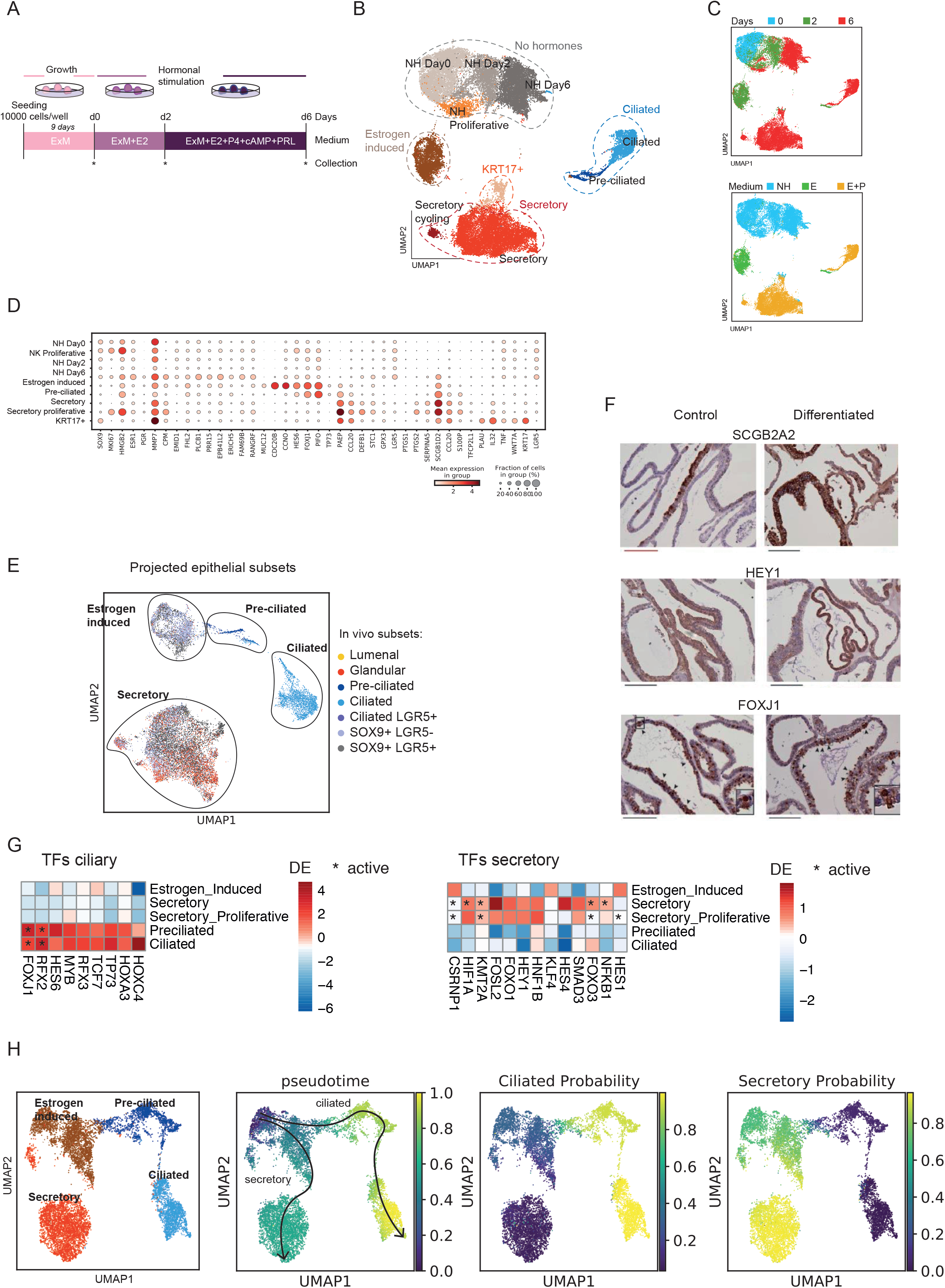
Endometrial organoids recapitulate epithelial responses to ovarian hormones. **(A)** Experimental timeline of endometrial organoid culture. Clonal organoids were derived in Expansion Medium (ExM) and then subjected to hormonal stimulation with estrogen (E2) followed by E2+progesterone (P4)+cyclic AMP (cAMP) and prolactin (PRL). The time points at which organoids were collected for scRNA-seq are shown with an asterisk. **(B)** UMAP (uniform manifold approximation and projection) projections of scRNA-seq data identifies major populations of cells. **(C)** UMAP representations coloured by days after the hormonal stimulation (top) and by the treatments (bottom). **(D)** Dot plot showing log_2_-transformed expression of selected genes that distinguish the main cell populations. **(E)** Predicted epithelial subsets of endometrial organoids using logistic classifier. Validation of SCGB2A2 and HEY1 (markers of secretory population) and combined staining for FOXJ1 and acetylated-a-tubulin with immunohistochemistry in undifferentiated and differentiated (hormonally-stimulated) organoids. Black arrowheads indicate ciliated cells with FOXJ1 positive nuclei. Nuclei are counterstained with hematoxylin. Scale bars = 250 μm (red), 200 μm (black. Representative images of three endometrial organoids from three different donors. **(G)** Heatmaps showing TFs differentially expressed in ciliated and secretory lineages, respectively. Color proportional to log-fold change, with asterisk (*) highlighting TFs whose targets are also differentially expressed (DE) (i.e. differentially activated TFs). **(H)** Cells able to respond to progesterone (PGR+ clusters) derived from clonal E001 individual and coloured by (from left to right): i) cluster labels as in Figure S7F; ii) Palantir pseudo-time; iii) Probability of cells to progress towards the ciliary lineage and, iv) Probability that the cell differentiates toward the secretory lineage. NH = No hormone; E = Estrogen; P = Progesterone; d = days.

To assess how similar the organoids are to their *in vivo* counterparts, we looked for specific markers of the clusters. For a more quantitative assessment, we projected the epithelial *in vivo* data onto the hormone-treated *in vitro* epithelial subsets. Assignments were made based on logistic regression predictions and cosine distances. The latter allowed us to identify genes that are correlated between both datasets. Prior to hormonal treatment, the majority of cells within the organoids are proliferative (*TOP2A, PCNA*) and all express the E2 receptor, *ESR1* (**Fig. 4B-D, Fig. S7A-B**). Two additional populations emerge when the organoids are treated with E2: (i) a estrogen-induced population expressing the progesterone receptor (*PGR*), a target gene of estrogen, and (ii) a pre-ciliated population sharing markers with the equivalent cluster defined *in vivo*. The best match for the estrogen-induced population is the *SOX9+LGR5−* population, while pre-ciliated organoid cells align with their pre-ciliated *in vivo* counterparts (**Fig. 4E; Fig. S7C-D and Table S8**). Upon further stimulation with P4, markers of more advanced stages of differentiation emerge in both secretory and ciliated populations. Secretory cells express secretory (*PAEP*, *DEFB1*) and glandular (*SCGB2A2*) markers. Ciliated cells express typical markers (*FOXJ1, TP73*) and closely match their *in vivo* counterparts. IHC confirmed the expression of glandular and ciliary markers on the endometrial organoids after stimulation (**Fig. 4F**). Glandular secretory epithelium is the best match for 26.54% (logistic regression) and 3.94% (cosine distances) of secretory cells *in vitro*, indicating organoids cells respond similarly to hormones as *in vivo* **(Fig. 4E, Table S8)**. The remaining cells in this cluster match to *SOX9* subsets, in keeping with continued proliferation of organoids in culture and less complete secretory differentiation.

To dissect the pathways driving hormonally-induced differentiation towards the ciliated and secretory lineages, we explored if TFs defining these lineages are similar *in vitro* and *in vivo* by looking at their differential expression and activity (**Fig. 4G, Table S9**). We calculated differentially expressed or active TFs in the hormonal substes *in vivo* and *in vitro*. WNT-activated TFs (*FOXJ1*) are present in the ciliated lineage whilst WNT-inhibitory TFs (*CSRNP1)* in the secretory lineage. NOTCH-induced TFs, *HEY1* and *HES1,* are activated in the secretory lineage. These results indicate that ovarian hormones activate similar pathways both *in vivo* and *in vitro*.

We therefore felt it is possible to reconstruct pseudotime and recapitulate cell fate decisions of epithelial cells. We performed scRNA-seq on two clonal organoids from one individual (E001) (**Fig. S7E**). Both clones show similar behaviour and were integrated under the same manifold (**Fig. S7F-G**). Annotation of clusters was performed based on known markers and clusters expressing progesterone receptors were selected to further reconstruct epithelial differentiation in response to hormones (**Fig. 4H, Fig. S7H**). A subset of cells emerging from the estrogen-induced population differentiates into pre-ciliated cells in response to estrogen and, following additional progesterone, into ciliated cells. The secretory lineage also emerges from the estrogen-induced population. Thus, there is a common progenitor for both lineages.

### WNT and NOTCH inhibition mediate epithelial differentiation

To test the role of WNT and NOTCH pathways in ciliated and secretory differentiation we cultured organoids in the presence of either inhibitors of NOTCH (DBZ or DAPT) or WNT (IWP-2 or XAV939). We used functional, histological and single-cell transcriptomics assays to assess the outcomes (**Fig. 5A**). Organoid viability is high under all conditions (**Fig. S8A**). scRNAseq analysis reveals a higher proportion of pre-ciliated and ciliated cells and lower proportion of secretory cells in the presence of DBZ (NOTCH inhibitor) (**Fig. 5B-D, Fig. S8B-C, Table S10**). This finding was validated by IHC and qPCR using two NOTCH inhibitors (DBZ and DAPT) (**Fig. 5E. Fig. S8D)**. Ciliated cells are virtually absent when WNT is inhibited, highlighting the strong dependence on this pathway for ciliary commitment, while the proportion of secretory cells is increased under these conditions (**Fig. 5D, Table S10**). The drive towards the secretory lineage under WNT inhibitory conditions (IWP-2 and XAV939) was further validated by qPCR and IHC (**Fig. 5E. Fig. S8D)**. These results provide more evidence that the balance between NOTCH and WNT signalling regulates commitment to secretory or ciliary lineages.

**Fig. 5.**
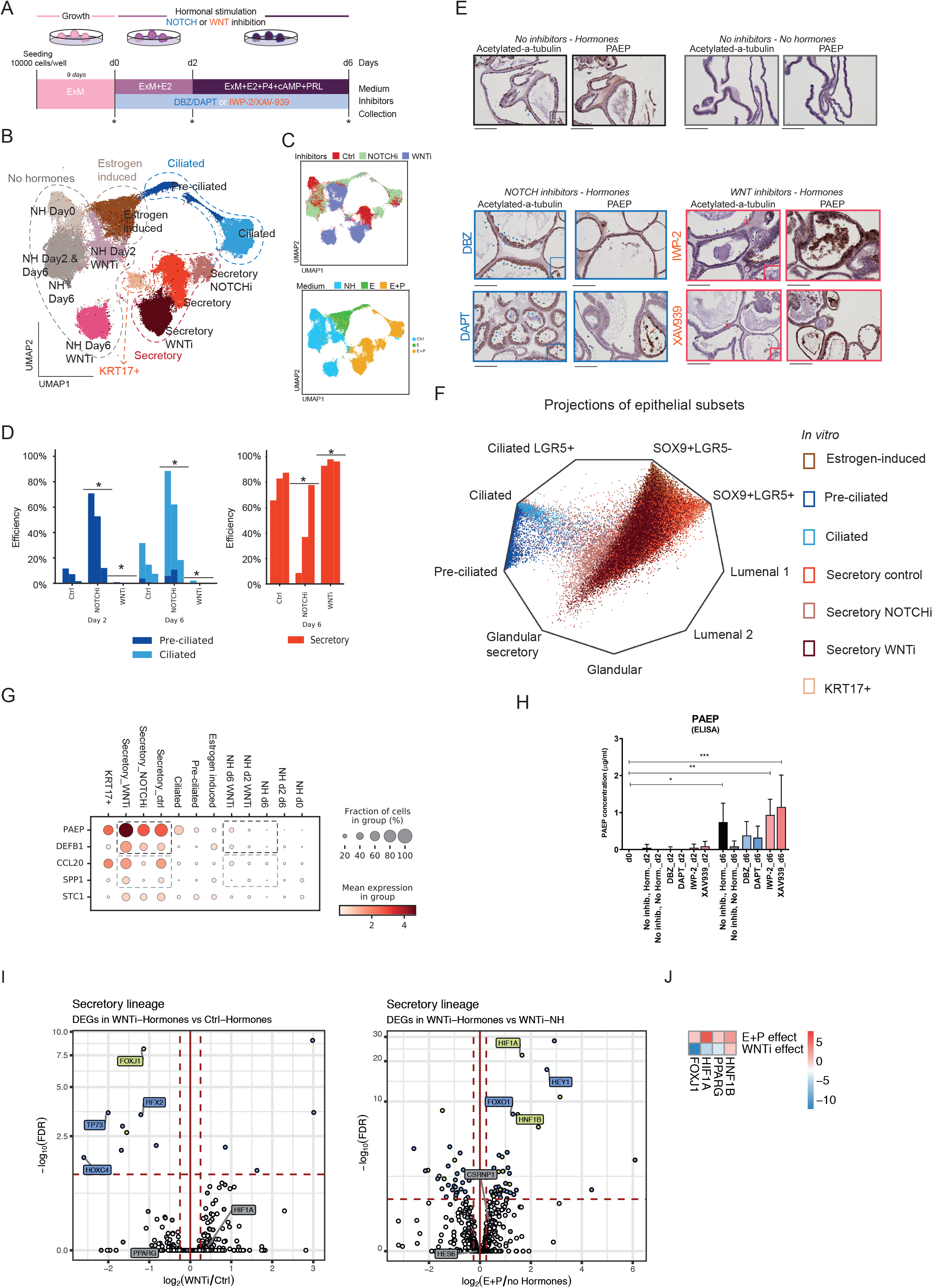
Signatures of WNT and NOTCH inform endometrial epithelial differentiation. **(A)** Experimental timeline of endometrial organoid culture. Organoids were treated with either NOTCH inhibitors (DBZ or DAPT) or WNT inhibitors (IWP-2 or XAV-939) upon initiation of hormonal stimulation. RSPO-1 was omitted from ExM in the presence of WNT inhibitors. Collection time points for scRNA-seq are shown with asterisks. **(B)** UMAP plots for scRNA-seq samples after WNT and NOTCH inhibition. **(C)** UMAP representations coloured by inhibitor treatments (top) and hormonal stimulation (bottom). **(D)** Bar plots showing enrichment of cells in ciliated and secretory clusters cells after NOTCH and WNT inhibition compared to untreated controls. The increased proportion of ciliated cells after NOTCH is inhibited and the decreased proportion after WNT inhibition are indicated with a star when statistically significant across the three genotypes using a paired t-test. **(E)** Immunohistochemistry for acetylated-a-tubulin (ciliary marker) and glycodelin (PAEP). Scale bars, 200 μm. Representative images of endometrial organoids derived from three different patients. **(F)** Radial representation of the cell type probabilities predicted by a logistic model trained on epithelial cells *in vivo*. The linear projection shows cells in each corner whenever a cell is predicted to belong to a given class with probability 1. **(G)** Dot plot showing the log_2_-transformed expression of genes characteristic of uterine secretions in epithelial subsets. **(H)** ELISA assay for glycodelin (PAEP) from supernatants of hormonally stimulated organoids treated with inhibitors, DBZ, DAPT, IWP-2 or XAV939. Shown are the mean with SD levels of expression relative to control conditions (no inhibitors but hormonally stimulated). Data from endometrial organoids from n=3 different donors. **(I)** Volcano plot representing differentially expressed genes in two comparisons within the secretory lineage: (i) analysis of effects of WNT inhibitor comparing cells cultured with and without WNT inhibitor. Progesterone is present in the media; (ii) analysis of effects of hormones with comparison of cells cultured with and without hormones. WNT inhibitor is present in the media. TFs that are significant in the *in vivo* dataset are highlighted. **(J)** Heatmap showing differential activities of TFs significant in the *in vivo* analysis. WNTi = WNT inhibitor; NOTCHi = NOTCH inhibitor; NH = No hormone; E = Estrogen; P = Progesterone.

Hormonal stimulation in the presence of WNT and NOTCH inhibitors modified the secretory cell transcriptome (**Fig. 5B**). To quantify the similarity of the new secretory populations with their *in vivo* counterparts, we measured changes in the overall transcriptome using cosine distances and logistic regression predictions. The computational projection of the *in vivo* dataset onto the *in vitro* dataset shows similar overlaps of the secretory populations emerging from WNT inhibitory conditions and controls with their *in vivo* counterparts (**Fig. 5F; Fig. S8E, Table S11**). Additionally, we measured the expression levels of genes encoding secretory products to check if this specific program is modified with NOTCH or WNT inhibition. Expression levels for *PAEP* and *DEFB1* are higher when WNT is inhibited and down-regulated with NOTCH inhibition (**Fig. 5G**). We validated the increase of one secretory product, PAEP, under WNT inhibitory conditions by ELISA (**Fig. 5H**). Although *PAEP* expression levels increase slightly in the presence of WNT inhibitors even in the absence of hormones, this is not significant (**Fig. 5G-H**).

We next dissected the regulatory programs in the secretory lineage by comparing expression and activity of TFs between populations emerging after treatment with and without hormones and WNT inhibition. The expression of NOTCH-regulated TFs (*HEY1*) is only upregulated in the presence of WNT inhibitors when hormones are present. This probably explains why WNT inhibitors are not sufficient to induce the secretory lineage on their own (**Fig. 5I-J, Table S12**). In the presence of hormones, WNT inhibition represses TFs characteristic of the ciliated lineage (*FOXJ1, TP73, RFX2*) (**Fig. 5I-J**). Switching off these genes may therefore drive secretory lineage differentiation.

## Discussion

Generating a cell atlas of the human uterus is essential to define the cell states and signalling pathways of the normal endometrium. This reference map, using samples from healthy women, will be important in understanding the molecular and cellular aberrations occurring in common conditions including infertility, endometriosis and endometrial carcinoma. The uterine lining in women of reproductive age is a challenging tissue to study due to difficulties accessing samples covering the dynamic changes occurring across all stages of the menstrual cycle. Here, we have used single-cell, spatial transcriptomics and high-resolution quantitative multiplex imaging to generate profiles of uterine cell states throughout the cycle (**Fig. 6A**). We focussed on the epithelial populations, as they are major players in endometrial function and pathology, confirmed by our comparison of gene signatures with genetic variants associated with endometrial disease. We utilise and develop novel computational tools to integrate and analyse scRNA-seq and spatial data, and investigate the molecular mechanisms driving epithelial differentiation in the glandular and lumenal microenvironments. We utilise our reference atlas to benchmark endometrial organoids, and engineer new lineage-specific endometrial epithelial organoids informed by signaling factors predicted by our *in vitro*/*in vivo* comparisons. Our work exemplifies the potential for leveraging human cell atlases as a blueprint for tissue engineering experiments.

**Fig. 6.**
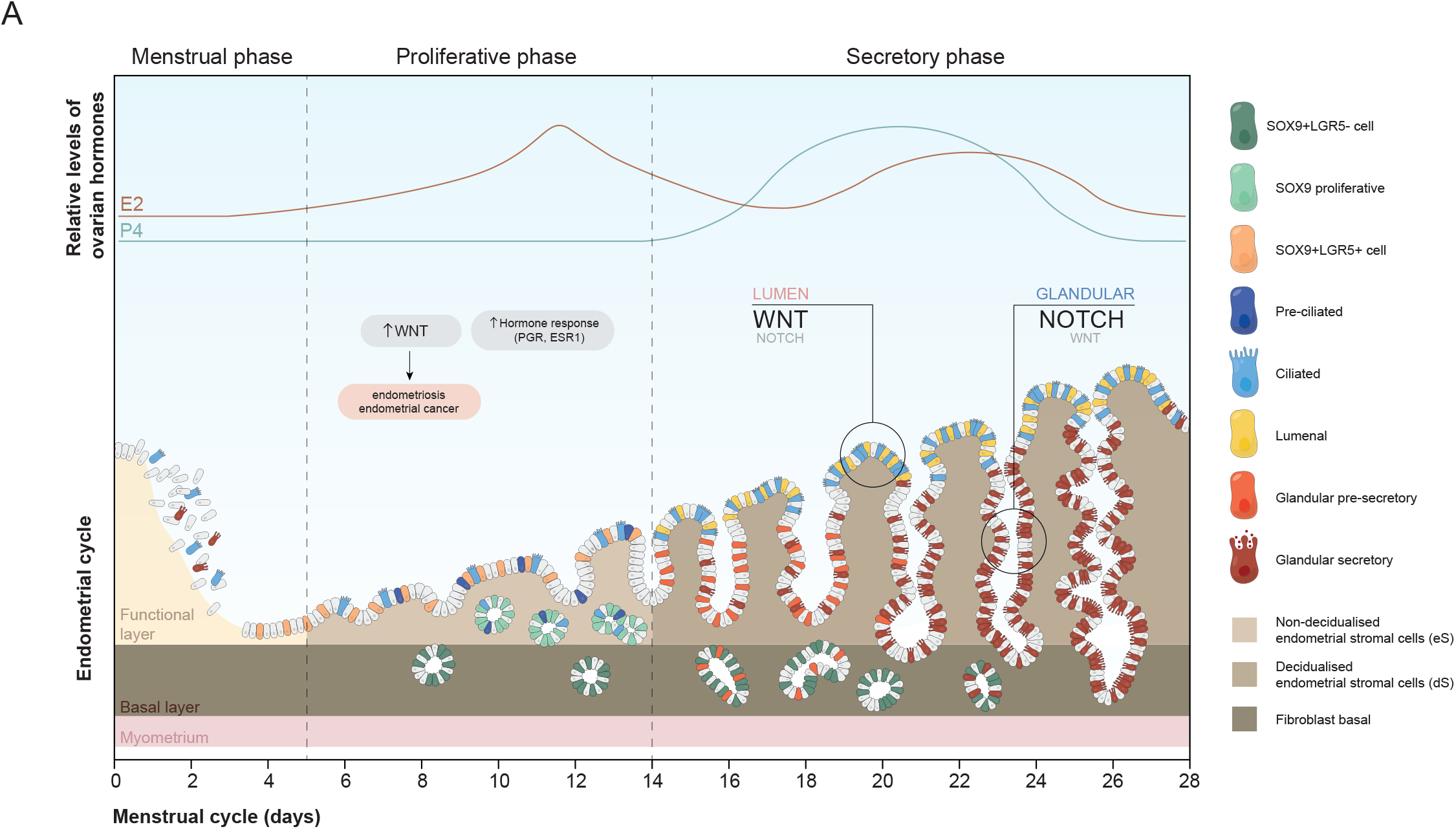
Proposed model of endometrial differentiation. **(A)** Schematic illustration showing our proposed model for temporal and spatial distribution of the epithelial and stromal subsets across the menstrual cycle. The proliferative phase is dominated by a WNT environment that promotes regeneration. Compartmentalisation of WNT and NOTCH signalling during the secretory phase promotes efficient differentiation towards the ciliated and secretory lineages. Epi = epithelium; uSMC = uterine smooth muscle cell; PV = perivascular; eS = non-decidualised endometrial stromal cells; dS = decidualised endometrial stromal cells.

In addition to our multi-omics approach, our uterine cell atlas has achieved three major advances in technologies and methods, which catalyse our discoveries. Firstly, we have accessed a diverse range of samples: uteri from transplant donors, hysterectomies and endometrial biopsies. This provides cells across the entire uterus including the endometrial basal layer and the myometrium. This led to the discovery of new cell states, including a novel population of fibroblasts (fibroblasts C7) restricted to the basal layer. Secondly, we sequence 48,867 cells in the poorly studied proliferative phase, revealing new cells such as a pre-ciliated population that appears in response to estrogen and is dependent on WNT signalling. Thirdly, our strategy of spatial mapping with 10x Genomics Visium and quantitative multiplex smFISH techniques means that we were able to determine the three-dimensional arrangements of cells described by suspension cell transcriptomics. By mapping cells into tissues with our new deconvolution method (*17*), we allocate epithelial cells into the three main endometrial layers: lumenal, functional and basal. Our newly expanded CellPhoneDB v3.0 analysis framework dissects the cell signalling mechanisms in the lumenal and glandular endometrial microenvironments. This reveals that NOTCH and WNT pathways control ciliated and secretory epithelial cell commitment.

Endometrial organoids are a powerful model for *in vitro* studies of the endometrial epithelium (*10*, *11*, *29*). We have systematically benchmarked the cellular composition of organoids relative to our *in vivo* reference map. Machine learning approaches, such as logistic regression scoring of expression profiles as well as correlation analysis, have been previously used to compare *in vitro* datasets with their *in vivo* counterparts (*30*, *31*). Using such an approach, we demonstrate that endometrial organoids recapitulate the *in vivo* response to hormones, thus providing compelling evidence for the validity of this model. While the ciliary lineage became fully differentiated, only ~30% of the secretory cells had a complete match with their *in vivo* counterparts. One possible explanation is that proliferation of organoids continues after treatment with progesterone, whereas uNK are the only endometrial cells that still divide *in vivo* after ovulation. We also compared TFs operating *in vivo* and *in vitro*, showing similar programs are induced. Further optimisation of the culture conditions to achieve more complete secretory differentiation is currently underway informed by these findings.

We envisage that our computational kit for *in vivo/in vitro* comparisons will be of general utility for tissue engineering experiments leveraging the Human Cell Atlas data as a blueprint.

Our extensive validation assessing hormonal responses of the endometrial organoids means that we could use them to test the effects of NOTCH and WNT signalling on epithelial cell fate (*10*). Inhibition of WNT signalling, which mimics the low WNT microenvironment in differentiated glands, inhibits ciliary commitment and induces secretory cells. In the presence of hormones these cells produce more secretions, likely through stronger silencing of ciliary genes. WNT inhibition alone does not result in secretory differentiation, as the NOTCH pathway is not induced without hormonal stimulation. These results reinforce previous findings suggesting tight coordination between these signalling pathways and ovarian hormones (*32*, *33*). NOTCH inhibitors promote the generation of ciliated cells as previously reported in fallopian tube (*34*) and endometrial organoids (*35*, *36*). We also show that secretory cells produced by NOTCH inhibition show lower expression of uterine milk molecules. Using single-cell mapping, we pinpoint the effect of NOTCH to early ciliary differentiation, as suggested by the strong effect NOTCH inhibition has on the novel pre-ciliated progenitor we have defined. Altogether, we demonstrate opposing roles of WNT and NOTCH in shaping distinct endometrial epithelial lineages, which *in vivo* is regulated by the boundaries set by the localisation of distinct cellular populations in the lumenal *versus* glandular microenvironments.

Our integrative map of cellular profiles of the normal endometrium will serve as an essential reference for studying many neglected endometrial disorders. Organoids, which can be bio-banked, have been established from samples of endometriosis and endometrial adenocarcinomas that resemble the original tumours (*10*, *11*, *37*, *38*). Our study shows that the combination of genomics, imaging and organoids can create a robust platform for studying endometrial physiology that will have a wide-ranging impact on women’s health and reproductive medicine.

## Acknowledgements

We gratefully acknowledge the Sanger Cellular Generation and Phenotyping (CGaP) Core Facility, and Sanger Core Sequencing pipeline for support with sample processing and sequencing library preparation. Anna Wilbrey-Clark for coordinating sequencing experiments; Antonio Garcia and Jana Eliasova for graphical images; Alison Kimber and Jill Riches for patient consent; Kay Elder and Tereza Cindrova-Davies for endometrial samples and derivation of organoid cultures; Martin Prete for web portal support; Graham Burton, Felipe Vieira Braga and Angela Goncalves for helpful discussions; Ni Huang and Alexander Aivazidis for data analysis; Stephen Williams and 10x R&D team for help on Visium analysis. The material from the deceased organ donor was provided by the Cambridge Biorepository for Translational Medicine. The endometrial biopsies were obtained from the Newcastle Uteroplacental Bank. We are grateful to the donors and donor families for granting access to the tissue samples.

## Author contributions

R.V-T and M.Y.T designed and supervised the experiments and analysis with contributions from S.A.T, O.B, L.G-A and K.N; R.H, C.S-S and R.V.T performed sampling and library prep with help from L.M, M.D, K.T.M, P.A, T.P, C.I.M, E.P and K.S-P; L.G-A, L-F.H and R.V-T analysed the sequencing data with contribution from K.P J.P, S.V.D, V.K, A.A; K.R, B.W, T.L, K.K, A.T, L.T, V.V and O. B performed and analysed the smFISH; L.G and C.L performed immunohistochemistry; K.N, R.F and M.Y.T performed the organoid experiments; L.M, M.J-L, L.C and A.M provided pathological expertise; R.V-T, A.M. and M.Y.T wrote the manuscript with contributions from L.G-A, K.R, K.N, L.M, M.R.S, O.B and S.A.T. All authors read and approved the manuscript.

## Data availability

Datasets can be accessed and downloaded through the web portals www.reproductivecellatlas.org (user: rca ; password: $Uj6mPXA). All codes used for data analysis are available from https://github.com/Ventolab/UHCA.

## Supplementary Materials

Materials and Methods

Fig S1 – S8

Table S1 – S18

References 39 – 59

