## Supplementary material for "Mapping the temporal and spatial dynamics of the human endometrium *in vivo* and *in vitro*"

**Affiliations:**
1 Wellcome Sanger Institute, Cambridge, UK
2 Centre for Trophoblast Research, University of Cambridge, Cambridge, UK
3 Department of Pathology, University of Cambridge, Cambridge, UK
4 EMBL-EBI, Wellcome Genome Campus, Hinxton, Cambridge, UK
5 Department of Surgery, University of Cambridge, Cambridge, UK
6 Cambridge NIHR Biomedical Research Centre, Cambridge, UK
7 Department of Pathology, Cambridge University Hospitals NHS Foundation Trust, Cambridge, UK
8 Department of Immunology, Genetics and Pathology and Science for Life Laboratory, Uppsala University, Uppsala, Sweden.
9 Newcastle University, Newcastle, UK
10 Theory of Condensed Matter Group, Cavendish Laboratory, University of Cambridge, JJ Thomson Ave, Cambridge CB3 0HE, UK

**This PDF includes:**

Materials and Methods
Fig S1 – S8
Table S1 – S18
References 39 – 59

**Material and Methods:**

**Uterine tissue retrieval**

*Whole uterine tissue from transplant organ donors*

Full thickness uterine wall samples were obtained from deceased transplant organ donors (A10, A13, A16, A30) after ethical approval (reference 15/EE/0152, East of England—Cambridge South Research Ethics Committee) and informed consent from the donor families. Uterus was removed within 1 hour of circulatory arrest.

Full thickness uterine wall samples were collected from four women during autopsy (Trv2, Trv3, Trv4 and Trv5). All four individuals died of non-cancer related causes traumatic injuries (Trv2, Trv3 and Trv4) and brain oedema (Trv5). Samples were collected within less than ten hours of death (post-mortem interval, PMI was 5, 2, 4 and 6 hours respectively. Once collected, all tissue biopsies were snap frozen in liquid nitrogen and subsequently stored at -80$^0$C. The use of these tissues was approved by the London, Surrey Research Ethics Committee (REC reference 17/LO/1801, 26/10/2017).

*Endometrial biopsies*

Endometrial biopsies were obtained from live donors with written informed consent from all participants from multiple centres:

i) Endometrial biopsies for sequencing were obtained from subjects recruited from the Newcastle Upon Tyne Hospitals after ethical approval (reference 16/NE/0167, North East - Newcastle & North Tyneside 1 Research Ethics Committee).

ii) Proliferative endometrial biopsy for deriving organoids were obtained from Addenbrooke's Hospital under ethical approval from the East of England-Cambridge South Research Ethics Committee (08/H0305/40).

iii) Endometrial scratch samples from secretory phase endometrium for deriving organoids were obtained from Bourne Hall Clinic under ethical approval from the East of England-Central Research Ethics Committee for the 'Biology of the Human Uterus in Pregnancy and Disease Tissue Bank' run by the Centre for Trophoblast Research (17/EE/0151).

Endometrial biopsies were obtained using a disposable endometrial cell sampler, starting from the uterine fundus and moving downward to the internal cervical ostium. None of the subjects were on hormonal treatments for at least 3 months prior to the procedure.

Endometrial tissues were staged based on standard histological criteria.

**Tissue processing**

All tissues for sequencing and spatial work were collected in saline (HypoThermosol biopreservation media) and stored at 4°C until processing. Tissue dissociation for all tissues was conducted within 24 hours of tissue retrieval in a two-step digestion protocol.

Step-1 Collagenase treatment: Tissue was transferred to a sterile 10 $mm^2$ tissue culture dish and cut into <1 $mm^3$ segments before being transferred to a 50 ml conical tube. Tissues were digested with 1 mg/ml collagenase type V in RPMI (Sigma-Aldrich), 0.1 mg/ml DNaseI (Sigma-Aldrich) supplemented with 10% (v/v) heat-inactivated fetal bovine serum (FBS; Gibco), 100 U/mL penicillin (Sigma-Aldrich) for 45 minutes at 37°C with intermittent shaking. Digested tissue was passed through a 100 µm filter, and cells collected by centrifugation (450×g for 5 minutes at 4°C). Cells were treated with 1X red blood cell (RBC) lysis buffer (eBioscience) for 5 minutes at room temperature and washed with flow buffer (PBS containing 5% (v/v) FBS and 2 mM EDTA) prior to cell counting. Protocol available at: dx.doi.org/10.17504/protocols.io.76thren

Step-2 Trypsin treatment: Pieces of tissue retained on the 100 µm filter were washed with PBS and further digested with Trypsin/EDTA 0.25% for 20 minutes at 37°C with intermittent shaking. Digested tissue was passed through a 100 µm filter, and cells collected by centrifugation (450×g for 5 minutes at 4°C). Cells were washed with flow buffer (PBS containing 5% (v/v) FBS and 2 mM EDTA) prior to cell counting. Protocol available at: dx.doi.org/10.17504/protocols.io.72dhqa6

**Tissue freezing**

Fresh tissue samples of human endometrium were embedded in cold OCT medium and flash frozen using a dry ice-isopentane slurry. Protocol available at: https://www.protocols.io/view/embedding-and-freezing-fresh-human-tissue-in-oct-u-95mh846

**Nuclei extraction**

Thick (200 µm) uterine sections were cryosectioned, dissected from OCT and kept in a tube on dry ice until subsequent processing. Nuclei were released *via* Dounce homogenisation as described previously (*39*).

**Endometrial organoid cultures**

Endometrial organoids were grown following the protocol described in (*10*). Briefly, organoids at 37°C in a humidified atmosphere of 5% $CO_2$. The medium was refreshed every 2-3 days and the organoids were passaged at an average ratio of 1:3 every 5-7 days. The organoid suspension was centrifuged for 6 minutes at 600×g between passaging steps. The passaged organoid pellet was resuspended in 25 µl ice cold Matrigel (Corning, 356231) droplets, plated in a 48-well plate (Costar, 3548), allowed to solidify at 37°C for 15-30 minutes, and covered with 250 µl endometrial organoid expansion medium (ExM). Components of ExM for culturing human endometrial organoids available at Table S18.

**Endometrial organoid dissociation**

Matrigel was removed using Cell Recovery Solution (Corning, 354253) for 1 hour on ice. Organoids were washed with cold PBS and broken up by 300 strokes of an automatic pipette (Eppendorf, Xplorer Plus 613-223), then incubated, first with pre-warmed accutase cell detachment solution (Corning, 25-058-CI) for 5 minutes at 37°C, and second with collagenase V (Sigma-Aldrich, C-9263) diluted in 10% FBS/Advanced DMEM/F12 for 15 minutes at 37°C. The digest was passed through a 40 µm nylon mesh cell strainer to purify a single-cell suspension. Where undigested fragments were present, the collagenase step was repeated. The final digest suspension was resuspended in ExM. Cells were diluted in trypan blue for live and dead cell counting using a haemocytometer.

**Hormonal stimulation and inhibition experiment of endometrial organoids**

Endometrial organoids were stimulated with hormones and treated with NOTCH γ-secretase inhibitors (DBZ, Tocris 4489 and DAPT, Tocris 2634) as well as WNT inhibitors (tankyrase inhibitor XAV939, Tocris 3748 and porcupine inhibitor IWP-2, Tocris 3533) for 6 days. First, 10,000 single cells were plated per 25 µL Matrigel droplet into a 48-well plate with ExM supplemented with Rho kinase inhibitor (Y-27632-CAS 146986-50-7) and CHIR 99021. Ten days after plating, organoids were primed with 10 nM E2 and treated with NOTCH and WNT inhibitors (20 µM DAPT, 1 µM DBZ, 2 µM IWP-2, 2 µM XAV939 in the ExM). Rspondin-1, a WNT signalling activator, was deprived from the ExM in the conditions where WNT inhibitors were used. After 48 hours, they were stimulated with 10 nM E2, 1 µM P4, 100 µg/ml cAMP, 20 ng/ml PRL) while still being treated with NOTCH and WNT inhibitors (20 µM DAPT, 1 µM DBZ, 2 µM IWP-2, 2 µM XAV939). Conditions in which: i) inhibitors but no hormones; ii) no inhibitors but hormones; and iii) no inhibitors and no hormones were added were used as controls.

**RNA extraction, cDNA synthesis and qRT-CR**

Total RNA was extracted using the RNeasy Micro Kit with on-column DNase treatment (Qiagen, 76004), following manufacturer's instructions. RNA was resuspended in 14 µl RNase-free water, and purity and concentration were determined using a NanoDrop 1000 spectrophotometer (Thermo Fisher Scientific). For cDNA synthesis, 500-1000 ng of total RNA was reverse transcribed using SuperScript VILO cDNA Synthesis Kit (Thermo Fisher Scientific, 11754050) following manufacturer's instructions. The extracted RNA was diluted into 5X VILO Reaction Mix containing random primers, dNTPs, and $MgCl_2$ and 10X SuperScript III Enzyme Blend containing SuperScript III Reverse Transcriptase, RNaseOUT Recombinant Ribonuclease Inhibitor, and proprietary helper protein. Reactions were then incubated for 10 minutes at 25°C, 1 hour at 42°C and 5 minutes at 85°C. A no-reverse transcriptase reaction was prepared for use as a control for genomic DNA contamination. Quantitative real-time PCR (qRT-PCR) was performed on a 7500HT Fast Real-Time PCR system (Applied Biosystems) using TaqMan Fast Advanced Master Mix (Thermo Fisher Scientific, 4444557) and Taqman gene-specific primer probes, following manufacturer's protocol. Initial denaturation was performed for 20 seconds at 95°C, followed by 40 amplification cycles of 3 seconds at 95°C and 30 seconds at 60°C. Each qRT-PCR reaction was performed in duplicate, alongside non-template controls. Expression levels were calculated using the comparative cycle threshold (Ct) method. Normalised expression levels were calculated as $2^{-\Delta Ct}$ where ΔCt = Ct (gene of interest) – Ct (geometric mean of housekeeping genes *HPRT1*, *TOP1*, and *TBP*). Statistical analyses were performed using GraphPad Prism 5.03 software. All data are represented as mean ± standard deviation (SD). Statistical difference between compared groups was evaluated by repeated measures one-way ANOVA for multiple groups compared to control treatments. A threshold of $p < 0.05$ was applied to indicate statistical significance.

Taqman probes used for RT-qPCR available at Table S15.

**Immunohistochemistry**

Endometrial tissue sections of proliferative and secretory phase, as well as endometrial organoid sections, were used for immunohistochemistry. Organoids were separated from Matrigel using the Cell Recovery Solution (Corning, 354253), washed with cold PBS, fixed in formalin (Sigma, F5554), embedded into 2% agarose (Melford, MB1200) and finally embedded into paraffin and sectioned at 4 µm thickness. The tissue and organoid sections were dewaxed with Histoclear (National Diagnostics, HS-200), cleared with 100% ethanol and rehydrated serially through 90%, 70%, 50% ethanol to PBS. Epitope retrieval was conducted in Access Revelation (AR) pH 6.4 (A.Menarini, MP-607-PG1) citrate buffer or Access Super (AS) pH 9 (A.Menarini, MP-606-PG1) Tris-EDTA buffer at 125°C in an Antigen Access pressure cooker (A.Menarini, MP-2008-CE). Sections were blocked with 2% serum (of the

species of the secondary antibody) in PBS, then incubated with primary antibody for 30 min at RT or overnight at 4°C followed by three washes with PBS. Sections were then incubated with biotin-conjugated secondary antibodies for 30 min at RT followed by three washes with PBS and subsequently incubated with Vectastain ABC-HRP reagent (Vector, PK-6100) for 30 minutes at RT followed by two washes with PBS. Stains were developed by applying di-aminobenzidine (DAB) substrate (Sigma, D4168) directly to the sections. Sections were counterstained with Carazzi's haematoxylin and mounted in glycerol/gelatin mounting medium (Sigma, GG1-10).
List of antibodies provided in Table S13.

**ELISA**

The concentration of the human placental protein 14 (PP14/PAEP/glycodelin) in the organoid culture supernatants was assayed using the RayBio Human PP14 ELISA kit (RayBiotech, ELH-PP14), following manufacturer's instructions. The samples were diluted 1:2 with assay diluent and plated in duplicate. For each sample, three biological replicates were used. The emitted absorbance at 450 nm was measured with Synergy HT microplate reader (BioTek). The mean absorbance for each set of duplicate standards and samples was calculated and the average zero standard optical density was subtracted. The standard concentration and absorbance values were used for the generation of a standard curve from which sample PP14 concentrations were extrapolated.

**Multiplexed smFISH and high-resolution imaging**

Large tissue section staining and fluorescent imaging was conducted largely as described previously (*40*). Sections were cut from FFPE blocks at a thickness of 5 µm using a microtome, placed onto SuperFrost Plus slides (VWR), and baked at 55°C to dry and ensure adhesion. Tissue sections were then processed using a Leica BOND RX to automate staining with the RNAscope Multiplex Fluorescent Reagent Kit v2 Assay (Advanced Cell Diagnostics, Bio-Techne), according to the manufacturers' instructions, in combination with immunohistochemistry (IHC) for EPCAM (*41*). Probes may be found in Table S14. Automated processing included baking at 60°C for 30 minutes and dewaxing, as well as heat-induced epitope retrieval at 95°C for 15 minutes in buffer ER2 and digestion with Protease III for 15 minutes. Tyramide signal amplification with Opal 520, Opal 570, and Opal 650 (Akoya Biosciences) was used to develop three RNAscope probe channels. IHC was conducted after RNAscope completion. A blocking step of 1 hour in Primary Antibody Diluent (Leica) was followed by rabbit anti-EPCAM (Abcam ab71916) at 1:1,500 at room temperature for 2 hours, and then HRP goat anti-rabbit IgG (Thermo G21234) at 1:1,500 at room temperature for 1 hour. Both antibodies were diluted in Primary Antibody Diluent. The IHC signal was developed using TSA-biotin (TSA Plus Biotin Kit, Perkin Elmer) and streptavidin-conjugated Atto 425 (Sigma Aldrich). Stained sections were imaged with a Perkin Elmer Opera Phenix High-Content Screening System, in confocal mode with 1 µm z-step size, using a 20× water-immersion objective (NA 0.16, 0.299 µm/pixel). Channels: DAPI (excitation 375 nm, emission 435-480 nm), Atto 425 (ex. 425 nm, em. 463-501 nm), Opal 520 (ex. 488 nm, em. 500-550 nm), Opal 570 (ex. 561 nm, em. 570-630 nm), Opal 650 (ex. 640 nm, em. 650-760 nm).

**10x Genomics Chromium GEX library preparation and sequencing**

Cells were loaded according to the manufacturer's protocol for the Chromium Single Cell 3' Kit v.2 or v.3.0 to attain between 2,000 and 10,000 cells per well. Library preparation was carried out according to the manufacturer's protocol. Individuals E001, B044 and B080 for the organoids experiments were multiplexed under the same reaction. Libraries were sequenced, aiming at a minimum coverage of 20,000 raw reads per cell, on the Illumina HiSeq 4000 or Novaseq 6000 systems; using the sequencing formats; read 1: 26 cycles; i7 index: 8 cycles, i5 index: 0 cycles; read 2: 98 cycles (3' Kit v.2) or read 1: 28 cycles; i7 index: 8 cycles, i5 index: 0 cycles; read 2: 91 cycles (3' Kit v.3).

**10x Genomics Visium library preparation and sequencing**

Ten micron cryosections were cut and placed in duplicate on Visium slides (beta product version). These were processed according to the manufacturer's instructions. Briefly, sections were fixed with cold methanol, stained with haematoxylin and eosin and imaged on a Hamamatsu NanoZoomer S60 before permeabilisation (endometrium 20 min), reverse transcription and cDNA synthesis using a template-switching protocol. Second-strand cDNA was liberated from the slide and single-indexed libraries prepared using a 10x Genomics PCR-based protocol. Libraries were sequenced (1 per lane on a HiSeq4000), aiming for 300M raw reads per sample with read lengths 28cy R1, 8cy i7 index, 0cy i5 index, 91cy read 2.

**Alignment, quantification and donor deconvolution of scRNA-seq data**

The libraries were mapped with STAR 2.7.3a, using a reference based on 10x Genomics' GRCh38 1.2.0 release (which was constructed from Ensembl 84, with gene_biotypes filtered to protein_coding, lincRNA, antisense and the various IG and TR genes and pseudogenes). While reads were required to align within exonic regions to be counted for single-cells samples, sample containing single-nuclei included all reads that aligned within whole transcript regions instead. STAR's STARsolo CB_UMI_Simple mode was used for the alignment and quantification, with the default soloCellFilter setting of 10x Genomics' "CellRanger2.2 3000 0.99 10" emulating CellRanger 2.2's cell/soup cutoff. A

BAM file of the alignment, sorted on coordinates, was also returned for downstream donor deconvolution for the organoid time course data set.

Donors within the multiplexed organoid samples (including E001, B044 and B080) were deconvolved using Souporcell (*42*). All three-donor samples were processed individually, and the resulting donor calls produced genotype variant features that allowed the association of donors identities across samples, using Souporcell's shared sample identification.

**Alignment, quantification and quality control of Visium data**

10X Genomics Visium sequencing data were aligned and quantified using the Space Ranger Software Suite (version 1.0.0, 10x Genomics Inc) using the GRCh38 human reference genome (official Cell Ranger reference, version 3.0.0). Spots were manually aligned to the paired H&E images by 10x Genomics. All spots under tissue (selection provided by 10x Genomics) were included in downstream analysis. The mean UMI counts per spot was 91283, and the mean number of genes per spot was 1962. Scanpy (*43*) (version 1.4.4) python package was used to load the spot-gene count matrix and perform downstream analysis. Custom plotting functions were implemented to produce paper figures (available at https://github.com/Ventolab/UHCA).

**Downstream scRNA-seq analysis**

*Doublet detection, alignment of data across different batches and clustering*

The 10X Genomics scRNA-seq data was analysed with Scanpy (*43*), with the pipeline largely following their recommended standard practices. In addition, we implemented a number of enhancements as described below.

Individual samples of single cell or single nuclei were initially analysed separately prior to be batch corrected into an integrated dataset, and had two-step diffusion doublet identification performed (*44*, *45*). In short, the initial doublet scoring was performed with Scrublet (*46*) on a per-sample basis, with the scores diffused by overclustering the cells and reporting each cluster's median value. Doublets were identified from a distribution of these scores centred at the median and using an MAD-derived standard deviation estimate, with statistically significant cells after FDR correction flagged as doublets. The second diffusion step takes place in a joint multi-sample manifold, with the frequency of identified doublets in granular (Leiden resolution 10) clusters serving as the basis for the distribution and the statistical significance analysis being repeated. The only deviation from the implementation described in (*45*) is a greater stringency in calling doublets, using Bonferroni for FDR correction, and a significance threshold of 0.01. This doublet detection method was applied to all samples; however, additional cells were identified as doublets in the organoid samples that were demultiplexed by souporcell(*42*), which identified doublets based on the presence of multiple genotype variant features found in single droplets.

After filtering cells with fewer than 500 genes and with more than 15% mitochondrial reads (20% for organoid samples), the samples were integrated using scVI (*47*). While the raw count matrices were used for single-cell, the expression of genes that were retrieved for single nuclei were denoised from ambient RNA prior to the manifold identification. For that task, decontX (*48*) was used on each sample separately. We then excluded genes associated with cell stage progression by excluding all marker genes for G2/M and S phase that are listed inside the Seurat package (*49*). The expression of the 5000 genes that were identified by scVI native method were modeled by its generative model with 64 latent variables for 500 iterations; however, the epithelial, endothelial and immune cells that were later identified in the *in vivo* dataset were subsequently reanalysed separately by using 16 latent variable instead, and also including all genes instead of the 5000 most variable genes. The resulting latent variables were used for neighbour identification, which is needed for Leiden clustering (*50*) and Uniform Manifold Approximation and Projection (UMAP) visualisation from the Scanpy package. The resulting clusters that appeared to be specific to a single donor, or that had lower numbers of gene expressed or lower percentage of mitochondrial expression were labeled and excluded from downstream analysis. In the epithelial and immune cell reanalysis, more cells were excluded there when they belonged to clusters that were exhibiting high doublet scores, and epithelial cells were subsampled to balance each donor contribution (see *"Projection of organoid data onto in vivo dataset").*

Code available at https://github.com/Ventolab/UHCA .

*Annotation of scRNA-seq datasets*

Identification, labeling and naming of the major cell types in the *in vivo* dataset was by manual inspection of marker genes and interpretation of these based on the literature. Marker genes are the genes differentially expressed across clusters using two approaches, where both evaluate the statistical significance of differences between cells belonging to a given cluster to all cells that do not. To further annotate the cell clusters in the *in vivo* dataset accounting for the *donor* effect, we first used DEseq2 (*51*) that would compare the cells were aggregated into *in silico* mini-bulks, which is the result of summing the raw expression of single cells separately according to their donor origin. Since some donors were absent in some cell type cluster due the menstrual cycle stage or the nature of the sample harvested, every pair of mini-bulk samples given to DESeq2 from a given donor (“cluster” and “rest” mini-bulks) there both excluded together if either of them were defined with less than 10 cells. As a second approach, the wilcoxon test was used to report genes that were differentially expressed; however, it accounted for change of sequencing depth instead. This was performed by partitioning cells into 4 groups corresponding to the quartiles of the sequencing depth of cells considered (independent of donors), and combining the 4 resulting Z-scores. In both cases, P-values were adjusted with the Benjamini and Hochberg method. Cluster cell identity was assigned manually using known marker genes from the literature.

For each cell, we estimated the cell cycle phase (G1, S or G2M) based on its expression of G2/M and S phase markers following the method described in (*52*) and implemented in (*52*) Scanpy score_genes_cell_cycle function. Briefly, the relative expression of these gene-sets was compared to the expression of a set of reference genes. Marker genes for G2/M and S phase were retrieved from the Seurat package.

*Efficiency of organoid differentiation*

To identify clusters predominantly appearing  organoid cultures upon treatment with WNT or NOTCH inhibitors, we evaluated the proportion of cells in each cluster coming from the organoids with and without inhibitors using Fisher’s exact test. Odds ratios were computed against the control samples (organoids grown with no inhibitors) at the matched time points and also separately for the 3 genotypes considered in the experiment. In addition, the robustness of the observed effect was evacuated by the comparison of cell fractions assigned to each cluster detailed by each genotype, using a paired t-test that compares these fractions of cells from samples that were treated with inhibitor to their respective control samples.

*Projection of organoid data onto in vivo dataset*

To identify transcriptomic similarities with the *in vivo* scRNA-seq epithelial subset, we used a regularized logistic regression approach. To best link variation in gene expression in organoids to changes that are similarly observed in donors for which we could infer the stage of the menstrual cycle, we subsampled the epithelial cells so that each donor could contribute to at most 1000 cells, which balances the cell number representing each stage that otherwise would be heavily dependant on the varying quality that is observed in the considered *in vivo* samples. In order to limit the influence of cell cycle on the projection results, we excluded a SOX9 proliferative cluster that was composed of a majority of cells that were identified to be in G2/M or S phase, and also excluded genes that would be used for the logistic model if there are among the G2/M and S markers genes from Seurat. Subsequently, we subset both datasets to their shared highly variable genes and further prune half of remaining genes that are not cell-type specific according to three heuristic measures (fold-increase, fold-increase x fraction-positive, fold-increase x fraction-positive$^{0.5}$) (*53*). Gene expression the log transformed and normalized by the maximum RNA expression for both *in vivo* cells and organoid cells. The model is trained with the epithelial cell types identified in the *in vivo* cells assigned identities, with 10,000 iterations. The model was used to classify cells found in the organoid cultures *in vivo* cells. In addition, a quantitative measure of the similarity was also reported by evaluating the cosine distance from single cells for the organoid cultures to the centroid expression defined by the cell type clusters identified in *in vivo* cells, where each vector compared either contains expression of all selected genes in a single cell or their respective mean expression for a given cell type cluster identified in the reference *in vivo* samples. As a mean to visualise the projection results, a radial projection was used as suggested by (*30*), where the position of cell overlaid corresponds to the weighted average of radially balanced unitary vectors (each pointing towards a different corner of a regular polygon), where the weight is either each posterior probability of cell to belong to given celltype by the logistic model, or transformed cosine distances. The latter transformation is a softmax function where each component is multiplied by 15 before being exponentiated, which allows highlighting the identity of cell type classes with highest cosine projection.

*Trajectory analysis of organoid data*

In order to identify the branching point where cells commit to a ciliated or secretory fate, we first identified cell clusters e in the organoid cultures. Then, we then used Palantir (*54*) on cells from the clusters that did exhibit a mean expression level of the progesterone receptor higher than 0.2, which in fact effectively excluded clusters for which that quantity was always less than 0.04. A randomly selected cell corresponding to proliferating cells at day 2 was selected as a cell of origin.

**Downstream analysis of 10x Genomics Visium data**

*Location of cell types in Visium data*

To spatially map cell types defined by scRNA-seq analysis within the Visium spatial transcriptomics data, we used our novel cell2location model (*17*). Briefly, the model decomposes multi-cell spatial transcriptomics data into cell type abundance estimates in a spatially-resolved manner. The model uses a hierarchical non-negative decomposition of the gene expression profiles at spatial locations (each with multiple cells) into the reference signatures. The reference signatures that are estimated from scRNA-seq profiles are an estimate of the average gene expression profiles for each cell cluster. Cell2location employs informative priors on the number of cells, located signatures, and average change in sensitivity between technologies. The parameters used are shown below.

First, the model derived expression signatures of cell types by calculating average expression counts of each gene in each cell type in the raw count scRNA-seq data, selecting genes expressed in at least 3 cells. Next, to obtain cell type locations, the model was trained on raw counts of Visium spatial data using the intersect of 19070 genes. Each Visium section was analyzed separately. The following model parameters were used (the remainder set to default values):

- **train_args**='n_iter': 30000, **posterior_args**='n_samples': 1000.
- **model_kwargs**= 'cell_number_prior': {'cells_per_spot': 8, 'factors_per_spot': 4}; 'gene_level_prior':{'mean': 1/2, 'sd': 1/4, 'mean_var_ratio': 1};

We visualise the absolute amount of mRNA contributed by each cell population to each spot. We used 5% percentile of the posterior distribution of this parameter (mRNA counts), representing the number of mRNA molecules confidently assigned to each cell type.

*Clustering of spots in Visium data*

Visium data was processed using Scanpy (*43*) following the tutorial on spatial transcriptomics data analysis (similar approach to thatne used for scRNA-seq analysis described above): normalization using a scaling factor of 10000; log-transformation; variable gene detection with 'Seurat' flavor; PCA; neighbourhood graph building and UMAP calculation. Each sample was analysed independently.

Clusters were defined by the Louvain algorithm (clustering resolution manually tuned) and assigned as myometrium or endometrium based on visual inspection of the H&E image of the tissue aligned with each spot. The proportions of mRNA derived from a cell cluster in the endometrium and myometrium were calculated by first averaging mRNA counts from a cell cluster predicted by cell2location across spots in each of the regions (endometrium or myometrium), and then normalising these values against the total mRNA contribution of the cell cluster to all spots in endometrium and myometrium. This was done for all cell clusters.

The cluster of spots corresponding to epithelial cells in the endometrium for sample 152807 (A30) was further clustered using Louvain. One of the spot subpopulations was excluded due to the low percentage of epithelial cells in the spot after visual inspection. The other "epithelial" spot subpopulations were labelled based on the endometrial layer they were found - epithelial basal, epithelial luminal and epithelial glandular. Epithelial layers were defined based on visual inspection of the spots displayed on top of the H&E image of the tissue.

Differentially expressed genes between each subpopulation of "epithelial" spots was calculated using the limma R package (*55*) taking each subpopulation and comparing it to all other cells.

**Calculating transcription factor activities in scRNA-seq and Visium data**

Transcription factor (TF) activities were estimated *via* the combined expression levels of their targets. Target genes were retrieved from Dorothea (*27*), a collection of TF-targets compiled from a range of different sources (*i.e*. manual curation, ChIP-Seq data and *in silico* predictions from TF binding motif scanning on gene-promoters and coexpression in tissue-specific transcriptional data), where TF-target relationships are scored from A to E according to their degree of confidence. Here we updated Dorothea regulons as follows: first, synonymous gene names were corrected; second, *bona fide* TF targets relationships manually curated from Uniprot were added as a new curated source (Table S16); third, signed and curated interactions were upgraded to score B; fourth, TRRUST (*56*) curated interactions were updated to v2_20180416 version and signed interactions supported by more than one pubmed were upgraded to A and; finally, we created a new category (AA) for the most trustable TF-target interactions that were either detected by all approaches or in more than 2 curated resources (Table S13-2). For each TF, we used the highest scored set of targets and required at least ten target genes, as in the original publication (*27*).

Next, we estimated Dorothea-modified TF activities by performing a GSEA-like analysis of the limma-derived gene expression signatures of each cluster (from both scRNA-seq and Visium datasets, respectively) with the *msVIPER* function in the Viper R package (*57*) version 1.22.0.

**<u>CellPhoneDB v3</u>**

To study the interactions between epithelial and other cell populations identified in our endometrial samples, we used an updated version of our CellPhoneDB approach (*28*). Briefly, we retrieved the interacting pairs of ligands and receptors satisfying the following criteria: 1) all the members were expressed in at least 10% of the cell clusters under consideration; and 2) at least one of the members in the ligand or the receptor is a differentially expressed gene. To account for the distinct temporal and spatial regulation of cells, we further classified the epithelial interactions based on i) phase of the menstrual cycle where cell subsets co-exist, as assigned by histological inspection by experts pathologists (see Table S1); ii) location of cells in the three main endometrial layers (luminal, glandular and basal) as assigned by unbiased clustering of visium spots and cell2location (*17*). In order to account for the complexity of WNT cell-cell signaling, several ligands and receptors functional as heteromeric complexes were further curated manually and re-annotated in the CellPhoneDB database (Table S17).

Code available at https://github.com/Ventolab/Cellphonedbv3

**Image analysis**

*Image stitching and manual annotation of select glands:*

Confocal image stacks were stitched as two-dimensional maximum intensity projections using the BIOP Perkin Elmer Acapella Stitcher (EPFL, Lausanne; https://www.perkinelmer.com/PDFs/downloads/TCH-Workflows-In-Depth-High-Content-Analysis-Operetta.pdf). Four 3-plex smFISH panels were analysed, with each image measuring at least 4mm in each direction.

*smFISH quantification:*

RNA spot quantification was achieved with a three step process:

1) Segmenting the glands within the tissue

Ilastik (*58*) was used to train a random-forest based pixel classifier to detect valid gland areas based on the Nuclear-DAPI channel and the Gland-EPCAM(IHC) channel. Three rounds of ilastik classification were used to achieve adequate rejection of off-target signal to only segment the glands.

2) Segmenting the RNA spots within the glands

Another ilastik pixel classifier was used on the spot channels to segment areas that corresponded to genuine spots that were situated in the glands segmented in step 1. The spots were verified by only including spots visible in one channel only. This was done to successfully remove blood inclusions, which gave a confounding signal across multiple channels.

3) The edge of the lumen was manually annotated using napari (*59*). The distance of each pixel on the image was calculated to the nearest point on the lumen edge. Then the total fluorescence intensity was measured for spots in glands and binned into intervals of distance away from the lumen. The gland area was also calculated for each distance interval. The spot fluorescence was divided by the gland interval to give a value of spot intensity that was normalised by area.

58. ilastik. *ilastik*, (available at https://www.ilastik.org/index.html).

59. napari. *napari*, (available at https://napari.org/).

Fig S1

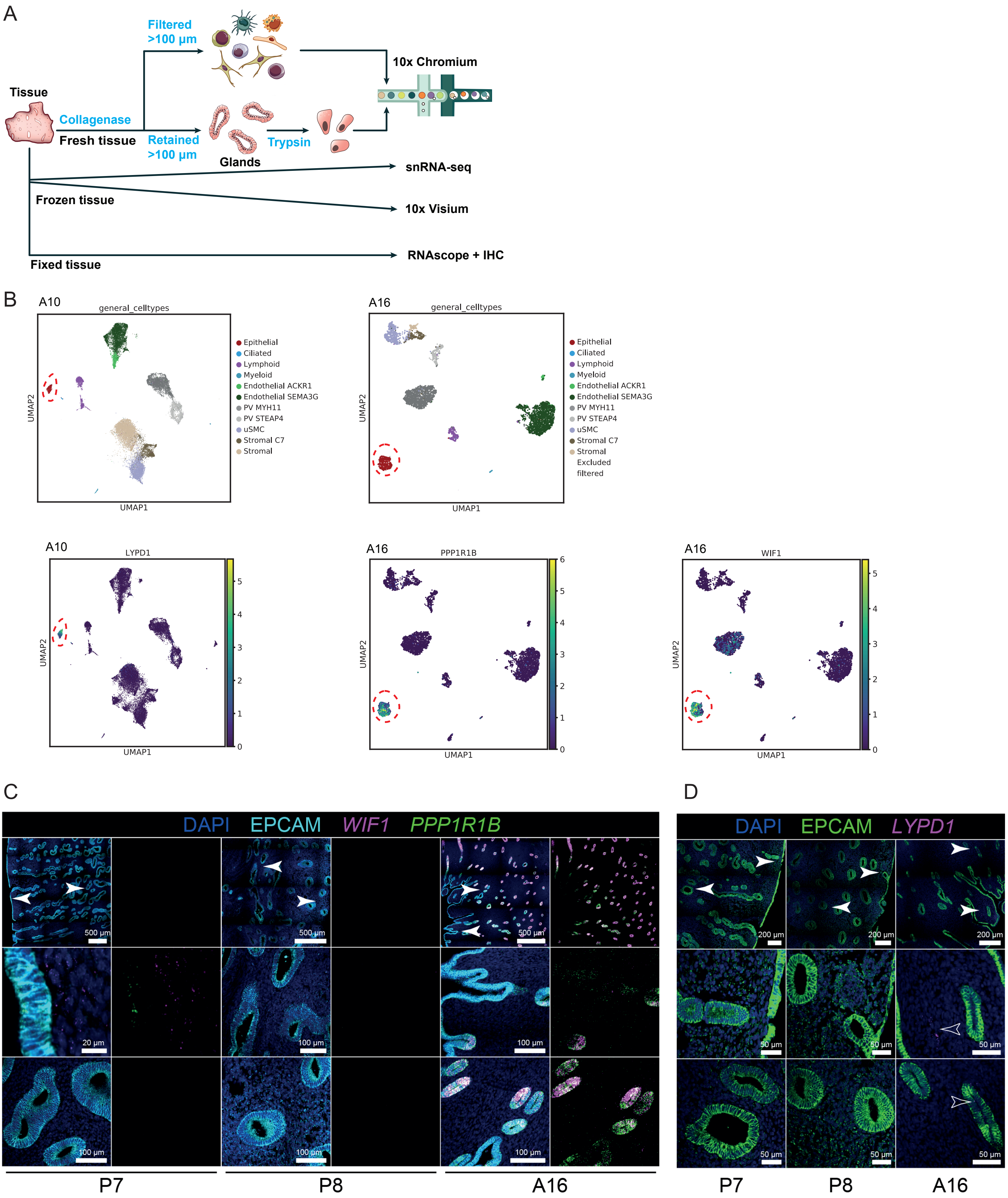
A
Filtered
>100 µm
10x Chromium
Tissue
Collagenase
Fresh tissue
Retained
>100 µm
Trypsin
Glands
snRNA-seq
Frozen tissue
10x Visium
RNAscope + IHC
Fixed tissue
B
A10
general_celltypes
A16
Epithelial
Ciliated
Lymphoid
Myeloid
Endothelial ACKR1
Endothelial SEMA3G
PV MYH11
PV STEAP4
uSMC
Stromal C7
Stromal
Excluded
filtered
UMAP1
UMAP2
LYPD1
PPP1R1B
WIF1
C
DAPI EPCAM WIF1 PPP1R1B
500 µm
20 µm
100 µm
P7
P8
A16
D
DAPI EPCAM LYPD1
200 µm
50 µm

**Fig. S1**

**Samples included in the study. (A)** Experimental workflow for the generation of cellular profiling of the uterus. In short, single-cell suspensions were obtained following two protocols: i) collagenase treatment to enrich for the stromal fraction, and ii) collagenase followed by trypsin to enrich for the glandular fraction. In addition, tissue blocks were processed for snRNA-seq and Visium experiments. **(B)** UMAP projections of scRNA-seq data for individuals A10 (left) and A16 (right). Both samples were discarded due to the presence of epithelial cells expressing markers that are not present in normal endometrium (*LYPD1* on A10 and *PPP1R1B* and *WIF1* on A16*)*. A10 had a clinical history of misscarriage (see Table S1). **(C)** High-resolution large-area imaging of uterine tissue sections, stained with smFISH for *PPP1R1B* and *WIF1*. These probes were exclusively highly expressed in the epithelial cells of A16. White arrowheads point areas shown at higher magnification. Scale bars shown on the bottom right of the image. **(D)** High-resolution large-area imaging of uterine tissue sections, stained with smFISH for *LYPD1.* These probes did not show expression for any of the tissues stained. No tissue for histology was collected from A10. White arrowheads point areas shown at higher magnification. Scale bars shown on the bottom right of the image.

Fig S2

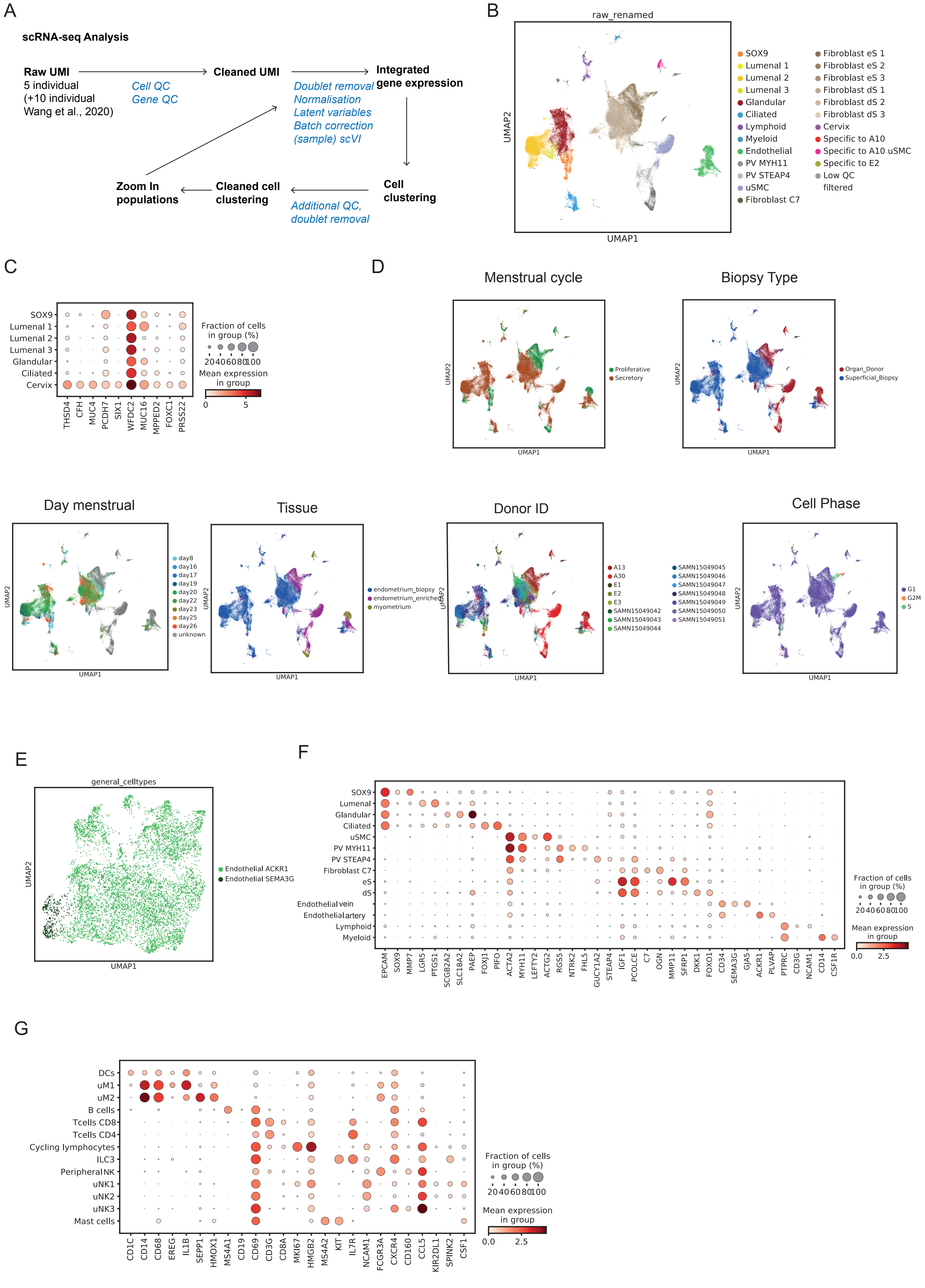
A
scRNA-seq Analysis
Raw UMI
5 individual
(+10 individual
Wang et al., 2020)
Cell QC
Gene QC
Cleaned UMI
Doublet removal
Normalisation
Latent variables
Batch correction
(sample) scVI
Integrated gene expression
Cell clustering
Additional QC, doublet removal
Cleaned cell clustering
Zoom In populations
B
raw_renamed
UMAP1
UMAP2
SOX9
Lumenal 1
Lumenal 2
Lumenal 3
Glandular
Ciliated
Lymphoid
Myeloid
Endothelial
PV MYH11
PV STEAP4
uSMC
Fibroblast C7
Fibroblast eS 1
Fibroblast eS 2
Fibroblast eS 3
Fibroblast dS 1
Fibroblast dS 2
Fibroblast dS 3
Cervix
Specific to A10
Specific to A10 uSMC
Specific to E2
Low QC filtered
C
SOX9
Lumenal 1
Lumenal 2
Lumenal 3
Glandular
Ciliated
Cervix
THSD4
CFH
MUC4
PCDH7
SIX1
WFDC2
MUC16
MPPED2
FOXC1
PRSS22
Fraction of cells in group (%)
20 40 60 80 100
Mean expression in group
0
5
D
Menstrual cycle
Proliferative
Secretory
Biopsy Type
Organ_Donor
Superficial_Biopsy
Day menstrual
day8
day16
day17
day19
day20
day22
day23
day25
day26
unknown
Tissue
endometrium_biopsy
endometrium_enriched
myometrium
Donor ID
A13
A30
E1
E2
E3
SAMN15049042
SAMN15049043
SAMN15049044
SAMN15049045
SAMN15049046
SAMN15049047
SAMN15049048
SAMN15049049
SAMN15049050
SAMN15049051
Cell Phase
G1
G2M
S
UMAP1
UMAP2
E
general_celltypes
Endothelial ACKR1
Endothelial SEMA3G
UMAP1
UMAP2
F
SOX9
Lumenal
Glandular
Ciliated
uSMC
PV MYH11
PV STEAP4
Fibroblast C7
eS
dS
Endothelial vein
Endothelial artery
Lymphoid
Myeloid
EPCAM
SOX9
MMP7
LGR5
PTGS1
SCGB2A2
SLC18A2
PAEP
FOXJ1
PIFO
ACTA2
MYH11
LEFTY2
ACTG2
RGS5
NTRK2
FHL5
GUCY1A2
STEAP4
IGF1
PCOLCE
C7
OGN
MMP11
SFRP1
DKK1
FOXO1
CD34
SEMA3G
GJA5
ACKR1
PLVAP
PTPRC
CD3G
NCAM1
CD14
CSF1R
Fraction of cells in group (%)
20 40 60 80 100
Mean expression in group
0.0
2.5
G
DCs
uM1
uM2
B cells
Tcells CD8
Tcells CD4
Cycling lymphocytes
ILC3
Peripheral NK
uNK1
uNK2
uNK3
Mast cells
CD1C
CD14
CD68
EREG
IL1B
SEPP1
HMOX1
MS4A1
CD19
CD69
CD3G
CD8A
MKI67
HMGB2
MS4A2
KIT
IL7R
NCAM1
FCGR3A
CXCR4
CD160
CCL5
KIR2DL1
SPINK2
CSF1
Fraction of cells in group (%)
20 40 60 80 100
Mean expression in group
0.0
2.5

**Fig. S2**

**Quality control of the scRNA-seq datasets. (A)** ScRNA-seq data analysis strategy (methods). In short, quality control was performed at the cell and gene level on the matrices generated by STARsolo. To integrate data from distinct individuals, data was batch corrected by each sample using scVI (*47*) on the 5000 most variable genes. The resulting latent variables were then used to define nearest neighbor associations needed by graph clustering that groups cells into clusters, which was performed using leiden graph from the Scanpy package. Clusters containing a high proportion of low-quality cells and doublets (defined by scrublet) were excluded. Re-clustering was performed on epithelial, endothelial and immune cells. **(B)** UMAP projections of scRNA-seq data from all tissue samples. Clusters corresponding to doublets, low QC cells and epithelial cells from the cervix were further excluded from the analysis. **(C)** Dot plot showing $log_2$-transformed expression of specific markers for the population labelled as "cervix", absent in organ donor samples (see (**Fig. S2D**)). Contamination from the cervix is possible due to the biopsy procedure (See Methods). **(D)** UMAP representations coloured by menstrual stage, biopsy type, menstrual day, tissue type, donor ID and cell cycle phase. Menstrual day was determined by LMP (see Table S1). **(E)** UMAP of sub-clustered endothelial populations. **(F)** Dot plot showing $log_2$-transformed expression of selected genes that distinguish the main cell populations. **(G)** Dot plot showing $log_2$-transformed expression of selected immune cell markers.

Fig S3

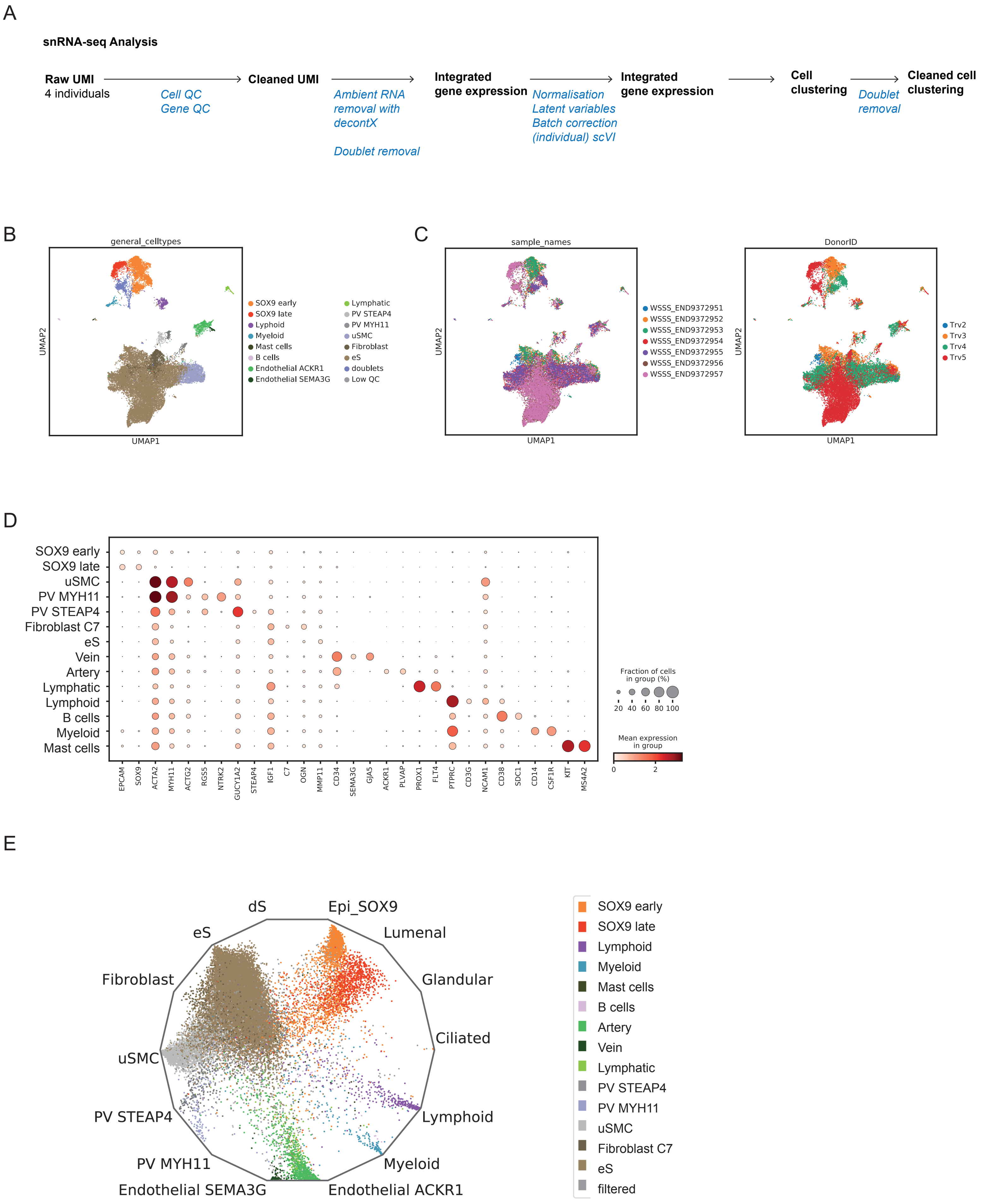
A
snRNA-seq Analysis
Raw UMI
4 individuals
Cell QC
Gene QC
Cleaned UMI
Ambient RNA removal with decontX
Doublet removal
Integrated gene expression
Normalisation
Latent variables
Batch correction (individual) scVI
Integrated gene expression
Cell clustering
Doublet removal
Cleaned cell clustering
B
general_celltypes
UMAP2
UMAP1
SOX9 early
SOX9 late
Lyphoid
Myeloid
Mast cells
B cells
Endothelial ACKR1
Endothelial SEMA3G
Lymphatic
PV STEAP4
PV MYH11
uSMC
Fibroblast
eS
doublets
Low QC
C
sample_names
UMAP2
UMAP1
WSSS_END9372951
WSSS_END9372952
WSSS_END9372953
WSSS_END9372954
WSSS_END9372955
WSSS_END9372956
WSSS_END9372957
DonorID
UMAP2
UMAP1
Trv2
Trv3
Trv4
Trv5
D
SOX9 early
SOX9 late
uSMC
PV MYH11
PV STEAP4
Fibroblast C7
eS
Vein
Artery
Lymphatic
Lymphoid
B cells
Myeloid
Mast cells
EPCAM
SOX9
ACTA2
MYH11
ACTG2
RGS5
NTRK2
GUCY1A2
STEAP4
IGF1
C7
OGN
MMP11
CD34
SEMA3G
GJA5
ACKR1
PLVAP
PROX1
FLT4
PTPRC
CD3G
NCAM1
CD38
SDC1
CD14
CSF1R
KIT
MS4A2
Fraction of cells in group (%)
20 40 60 80 100
Mean expression in group
0 2
E
dS
Epi_SOX9
eS
Lumenal
Fibroblast
Glandular
Ciliated
uSMC
PV STEAP4
Lymphoid
PV MYH11
Myeloid
Endothelial SEMA3G
Endothelial ACKR1
SOX9 early
SOX9 late
Lymphoid
Myeloid
Mast cells
B cells
Artery
Vein
Lymphatic
PV STEAP4
PV MYH11
uSMC
Fibroblast C7
eS
filtered

**Fig. S3**

**Quality control of the snRNA-seq datasets. (A)** SnRNA-seq data analysis strategy (methods) (see Figure S2A). Prior to data integration, ambient RNA was removed with decontX (*48*). **(B)** UMAP projections of snRNA-seq data from all tissue samples. Clusters corresponding to doublets and low QC cells were further excluded from the analysis. **(C)** UMAP representations coloured by donor ID and sample ID. **(D)** Dot plot showing $log_2$-transformed expression of selected genes that distinguish the main cell populations. **(E)** Radial representation of the cosine distances similarity for single cells obtained from snRNA-seq to the centroids of cell types defined by scRNA-seq.

Fig S4

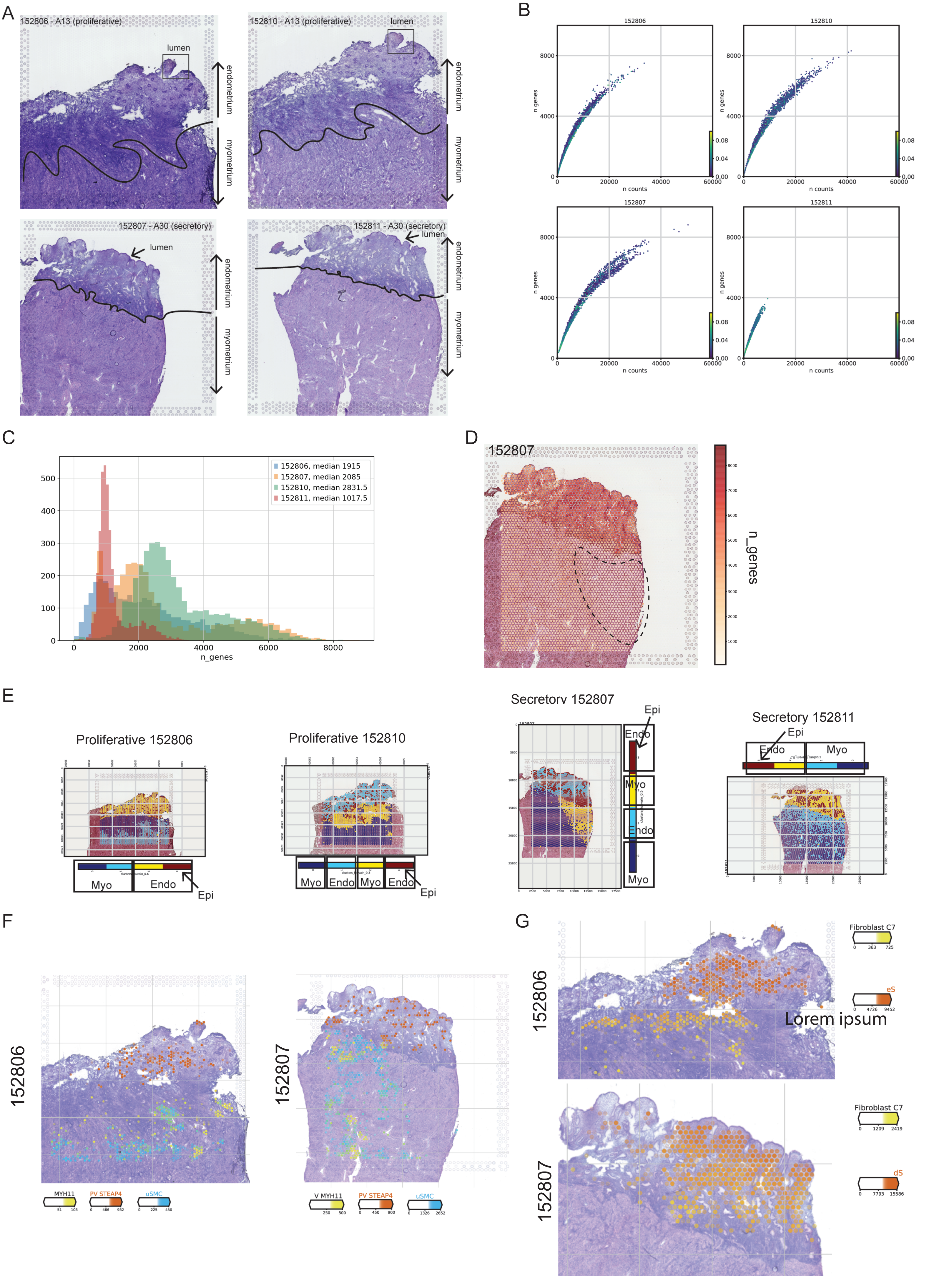

A
152806 - A13 (proliferative)
152810 - A13 (proliferative)
152807 - A30 (secretory)
152811 - A30 (secretory)
lumen
endometrium
myometrium
B
152806
152810
152807
152811
n genes
n counts
C
152806, median 1915
152807, median 2085
152810, median 2831.5
152811, median 1017.5
n_genes
D
152807
n_genes
E
Proliferative 152806
Proliferative 152810
Secretory 152807
Secretory 152811
Myo
Endo
Epi
F
152806
152807
MYH11
PV STEAP4
uSMC
G
152806
152807
Fibroblast C7
eS
dS
Lorem ipsum

**Fig. S4**

**Quality control of the Visium slides. (A)** Haematoxylin and eosin staining of the slides in the Visium arrays. Four individuals were selected: A13 was in their proliferative phase and A30 in the secretory phase. Two sections 100 µm apart were analysed. Lumenal epithelium was well preserved in individual A30 and in a small region of A13. **(B)** Scatter plots show the number of genes over the number of counts, where each dot is a feature of the Visium slide. Plots are coloured by the percentage of mitochondrial genes. **(C)** Bar plots showing number of genes on each of the samples. A bimodal distribution corresponding to endometrium and myometrium was shown on each of the cases. **(D)** Visualisation of the number of genes on the Visium slides of sample A30 (152807 slide). A zone with low quality is highlighted in the image. This is probably caused by a technical artifact. No pattern like this was seen in other samples. **(E)** Unbiased clustering of Visium spots defined by Louvain algorithm. (**F)** Estimated amount of mRNA (colour intensity) contributed by each cell population to each spot (colour) shown over the H&E image of the proliferative (sample 152806, A13) and secretory (sample 152807, A30) endometrium. Endo = Endometrium; Myo = Myometrium; Epi = Epithelial; uSMC= uterine smooth muscle cell; PV= perivascular.

Fig S5

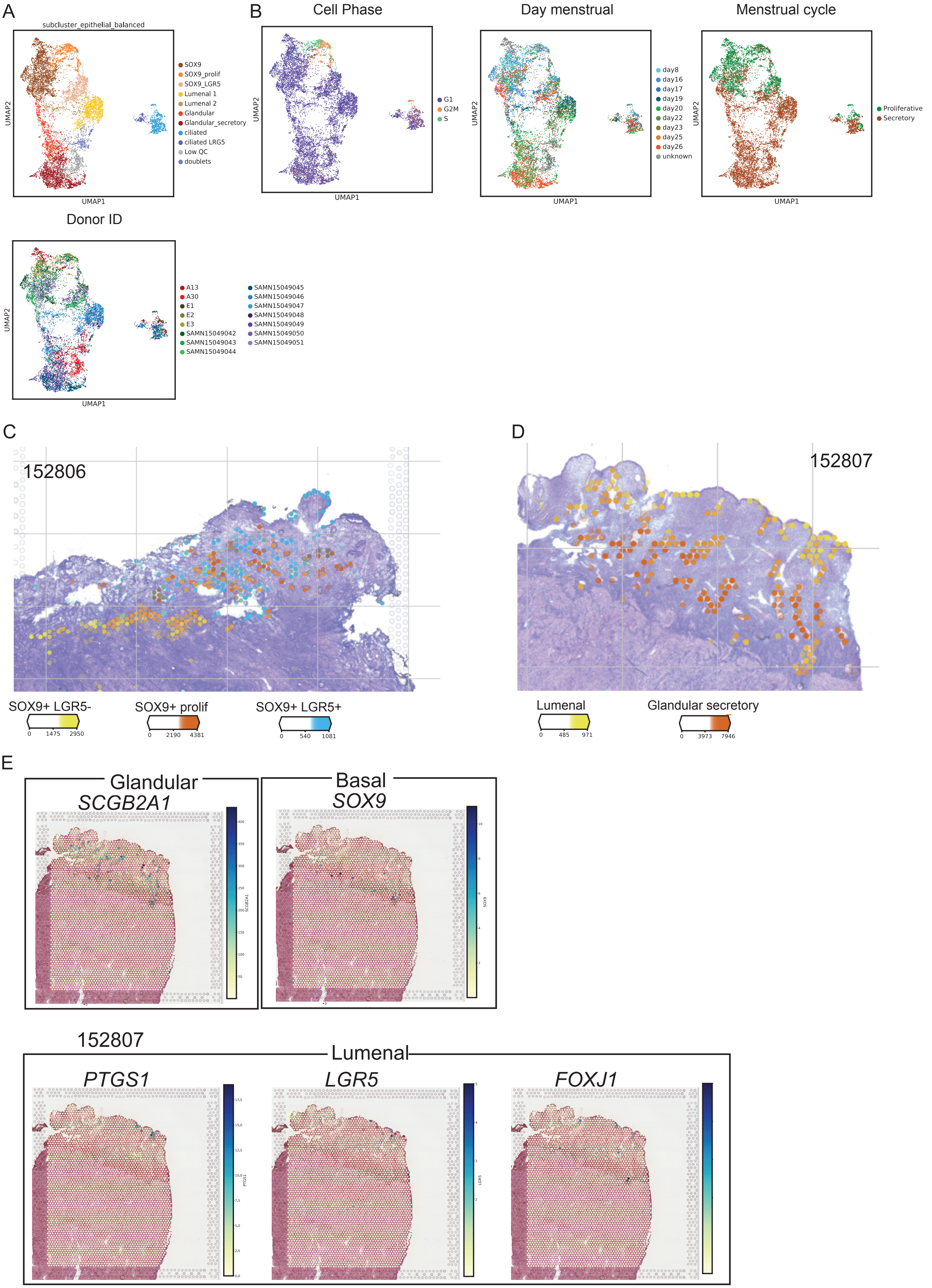
A
subcluster_epithelial_balanced
SOX9
SOX9_prolif
SOX9_LGR5
Lumenal 1
Lumenal 2
Glandular
Glandular_secretory
ciliated
ciliated LRG5
Low QC
doublets
UMAP1
UMAP2
B
Cell Phase
G1
G2M
S
Day menstrual
day8
day16
day17
day19
day20
day22
day23
day25
day26
unknown
Menstrual cycle
Proliferative
Secretory
Donor ID
A13
A30
E1
E2
E3
SAMN15049042
SAMN15049043
SAMN15049044
SAMN15049045
SAMN15049046
SAMN15049047
SAMN15049048
SAMN15049049
SAMN15049050
SAMN15049051
C
152806
SOX9+ LGR5-
0
1475
2950
SOX9+ prolif
0
2190
4381
SOX9+ LGR5+
0
540
1081
D
152807
Lumenal
0
485
971
Glandular secretory
0
3973
7946
E
Glandular
SCGB2A1
Basal
SOX9
152807
Lumenal
PTGS1
LGR5
FOXJ1

**Fig. S5**

**Spatio-temporal regulation of epithelial cells. (A)** UMAP projections of scRNA-seq data from epithelial cells. We performed a donor-balanced subsampling (1000 cells maximum for each donor). Clusters corresponding to doublets and low QC cells were further excluded from the analysis. **(B)** UMAP representations coloured assigned by cell phase, donor, menstrual stage and day of the menstrual cycle. Menstrual day was determined by LMP (see Table S1). **(C)** Number of mRNA molecules per spot (colour intensity) confidently assigned to each epithelial subpopulation (colour) in the proliferative phase (A13 - 152806). **(D)** Number of mRNA molecules per spot (colour intensity) confidently assigned to each epithelial subpopulation (colour) in the secretory phase (A30 - 152807). **(E)** Estimated proportion of mRNA coming from epithelial subsets in the early-proliferative phase (A30, 152807 slide).

Fig S6

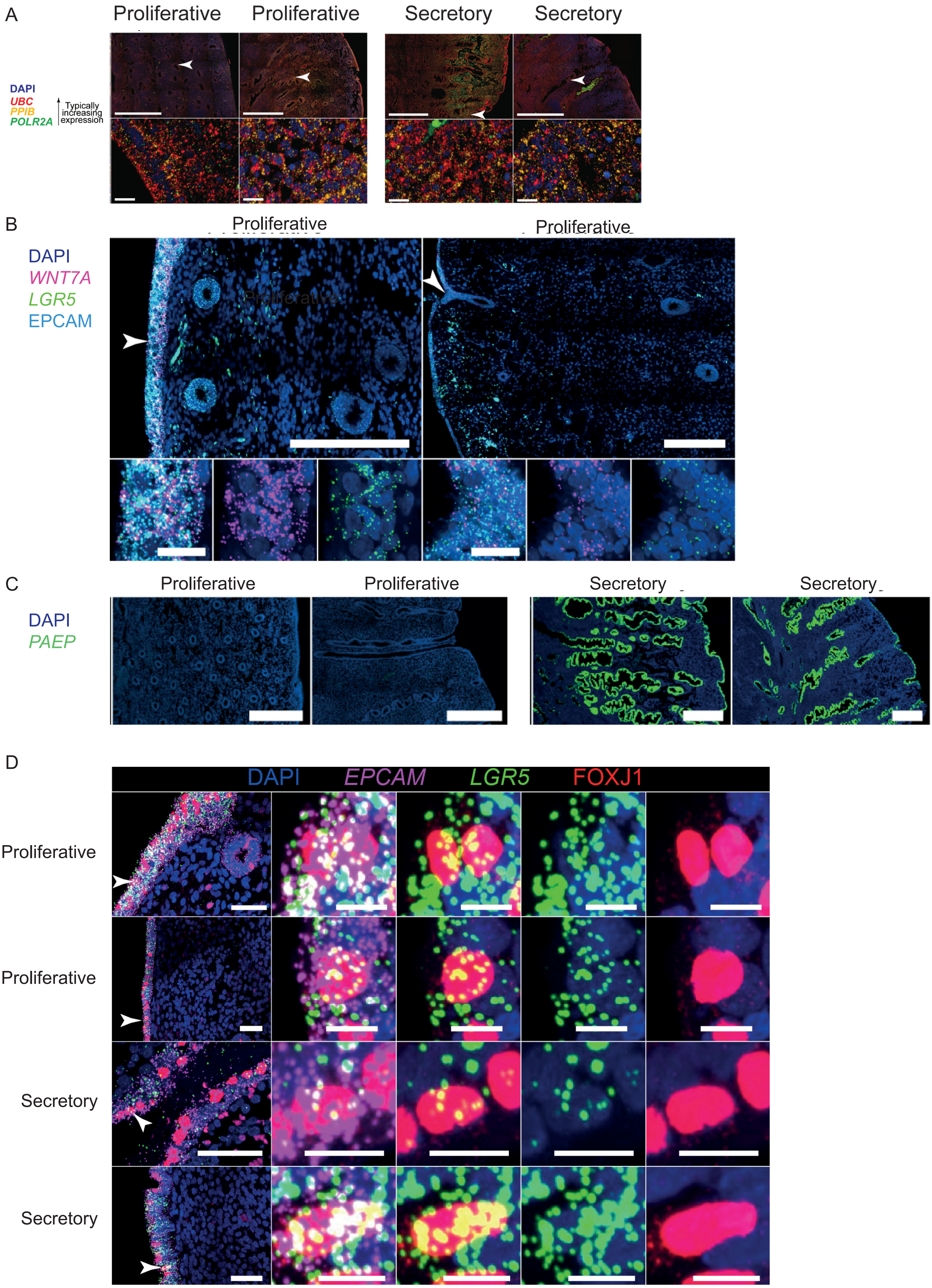
A
Proliferative
Proliferative
Secretory
Secretory
DAPI
UBC
PPIB
POLR2A
Typically increasing expression
B
Proliferative
Proliferative
DAPI
WNT7A
LGR5
EPCAM
C
Proliferative
Proliferative
Secretory
Secretory
DAPI
PAEP
D
DAPI
EPCAM
LGR5
FOXJ1
Proliferative
Proliferative
Secretory
Secretory

**Fig. S6**

**Spatially-resolved single-cell transcriptomic expression of proliferative markers by smFISH. (A)** Molecular integrity of uterine tissues was validated by multiplexed smFISH staining of sections for constitutively expressed genes of different typical expression levels (*UBC* - high; *PPIB* - moderate; *POLR2A* - low), which demonstrated strong signals irrespective of sample or region. White arrowheads indicate epithelial regions shown at higher magnification (bottom). Top scale bars = 1 mm; other scale bars = 20 µm. Representative images of four proliferative and four secretory endometrial samples from eight different donors. **(B)** High-resolution large-area imaging of two uterine tissue sections in the proliferative phase, stained with smFISH for *WNT7A* and *LGR5* (*SOX9+LGR5+* epithelial markers). White arrowheads indicate lumenal region shown at higher magnification (bottom). Representative images of four proliferative endometrial samples from four different donors. Top scale bars = 250 uM, bottom scale bars = 25 uM. **(C)** High-resolution large-area imaging of four endometrial sections stained with *PAEP*. Scale bars = 1 mm. Representative images of four proliferative and four secretory samples from eight different donors. **(D)** High-resolution large-area imaging of uterine tissue sections stained with smFISH for *EPCAM* and *LGR5*, combined with protein staining of FOXJ1. White arrowheads indicate cells with dual FOXJ1 and *LGR5* staining. Scale bars, left = 50 µm, other = 10 µm.

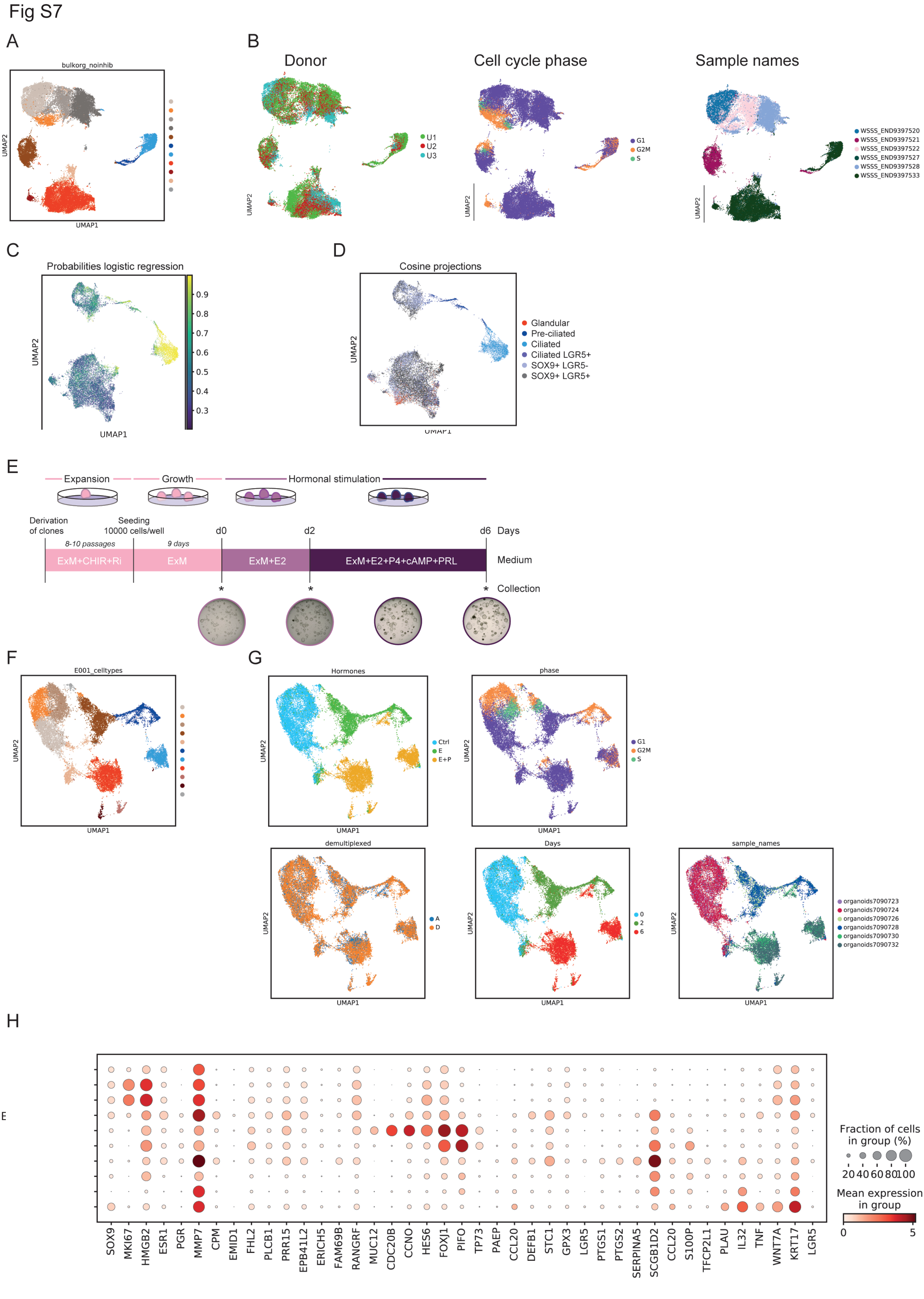
Fig S7
A
bulkorg_noinhib
UMAP1
UMAP2
B
Donor
U1
U2
U3
Cell cycle phase
G1
G2M
S
Sample names
WSSS_END9397520
WSSS_END9397521
WSSS_END9397522
WSSS_END9397527
WSSS_END9397528
WSSS_END9397533
C
Probabilities logistic regression
D
Cosine projections
Glandular
Pre-ciliated
Ciliated
Ciliated LGR5+
SOX9+ LGR5-
SOX9+ LGR5+
E
Expansion
Growth
Hormonal stimulation
Derivation of clones
Seeding 10000 cells/well
d0
d2
d6
Days
8-10 passages
9 days
ExM+CHIR+Ri
ExM
ExM+E2
ExM+E2+P4+cAMP+PRL
Medium
Collection
F
E001_celltypes
G
Hormones
Ctrl
E
E+P
phase
demultiplexed
A
D
Days
0
2
6
sample_names
organoids7090723
organoids7090724
organoids7090726
organoids7090728
organoids7090730
organoids7090732
H
Fraction of cells in group (%)
20 40 60 80 100
Mean expression in group
0
5
SOX9 MKI67 HMGB2 ESR1 PGR MMP7 CPM EMID1 FHL2 PLCB1 PRR15 EPB41L2 ERICH5 FAM69B RANGRF MUC12 CDC20B CCNO HES6 FOXJ1 PIFO TP73 PAEP CCL20 DEFB1 STC1 GPX3 LGR5 PTGS1 PTGS2 SERPINA5 SCGB1D2 CCL20 S100P TFCP2L1 PLAU IL32 TNF WNT7A KRT17 LGR5

**Fig. S7**

**Quality control of organoid scRNA-seq dataset. (A)** UMAP projections of scRNA-seq data from all organoid samples. **(B)** UMAP representations coloured by donor ID. The three donors correspond to E001, B044 and B080. **(C)** Logistic regression probabilities. **(D)** Cosine distances projections. **(E)** Experimental timeline of endometrial organoid culture. Clonal organoids were derived in Expansion Medium (ExM) with CHIR99021 (CHIR) and ROCK inhibitor Y-22763 (Ri), grown in ExM and then subjected to hormonal stimulation with estrogen (E2) followed by E2+progesterone (P4)+cyclic AMP (cAMP) and prolactin (PRL). The time points at which organoids were collected for scRNA-seq are shown with an asterisk. Representative bright field images of organoids for some of the timepoints are shown. **(F)** UMAP projections of scRNA-seq data from two clonal organoids derived from individual E001. **(G)** UMAP representations coloured by hormonal stimulation, cell cycle phase, individual clone, days after hormonal stimulation, sample ID. **(H)** Dot plot showing $log_2$-transformed expression of selected genes that distinguish the main cell populations.

Fig S8

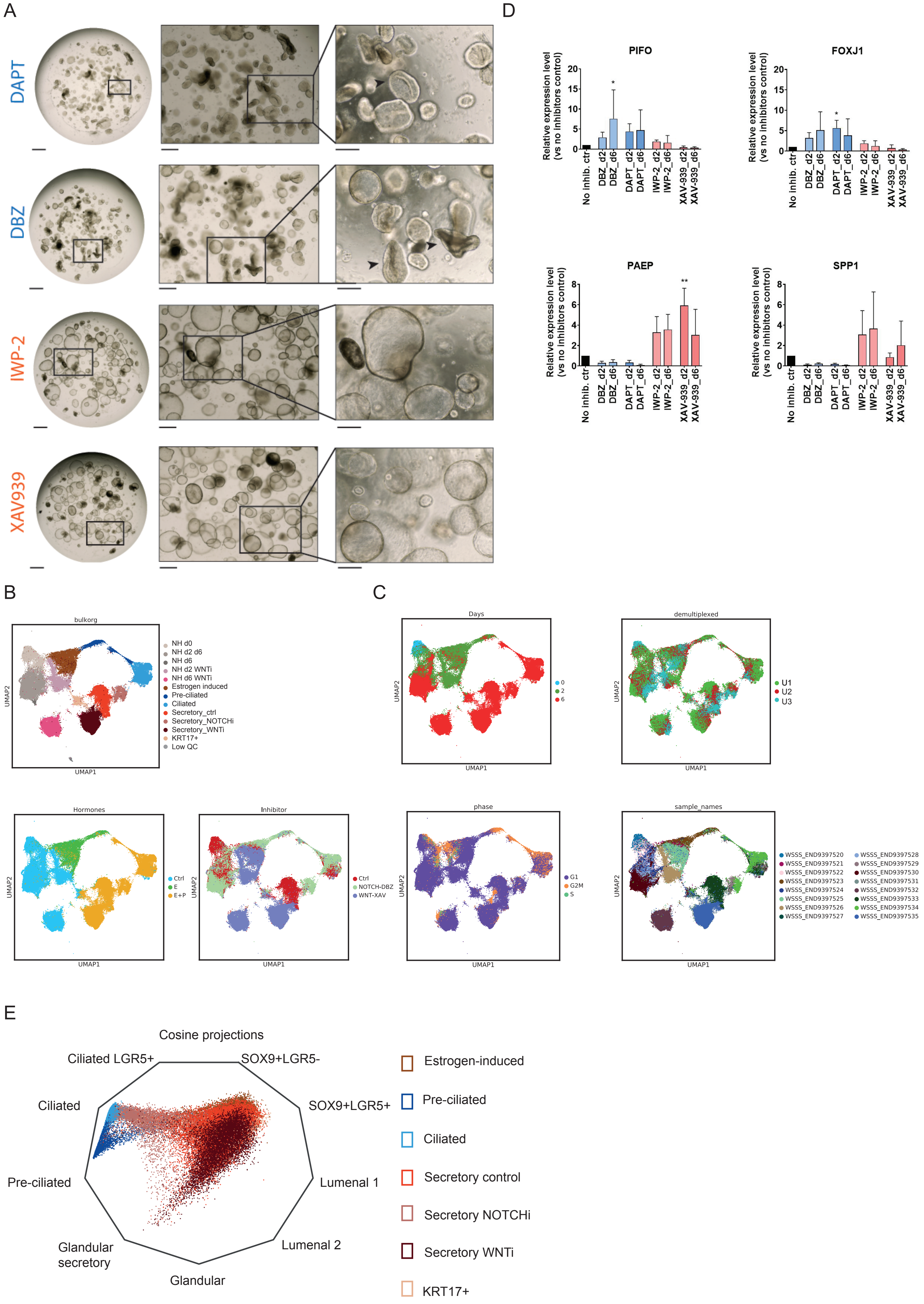
A
DAPT
DBZ
IWP-2
XAV939
D
PIFO
FOXJ1
PAEP
SPP1
Relative expression level (vs no inhibitors control)
No inhib. ctr
DBZ_d2
DBZ_d6
DAPT_d2
DAPT_d6
IWP-2_d2
IWP-2_d6
XAV-939_d2
XAV-939_d6
B
bulkorg
NH d0
NH d2 d6
NH d6
NH d2 WNTi
NH d6 WNTi
Estrogen induced
Pre-ciliated
Ciliated
Secretory_ctrl
Secretory_NOTCHi
Secretory_WNTi
KRT17+
Low QC
UMAP1
UMAP2
Hormones
Ctrl
E
E+P
Inhibitor
Ctrl
NOTCH-DBZ
WNT-XAV
C
Days
0
2
6
demultiplexed
U1
U2
U3
phase
G1
G2M
S
sample_names
WSSS_END9397520
WSSS_END9397521
WSSS_END9397522
WSSS_END9397523
WSSS_END9397524
WSSS_END9397525
WSSS_END9397526
WSSS_END9397527
WSSS_END9397528
WSSS_END9397529
WSSS_END9397530
WSSS_END9397531
WSSS_END9397532
WSSS_END9397533
WSSS_END9397534
WSSS_END9397535
E
Cosine projections
Ciliated LGR5+
SOX9+LGR5-
SOX9+LGR5+
Ciliated
Pre-ciliated
Lumenal 1
Lumenal 2
Glandular secretory
Glandular
Estrogen-induced
Pre-ciliated
Ciliated
Secretory control
Secretory NOTCHi
Secretory WNTi
KRT17+

**Fig. S8**

**NOTCH and WNT inhibition. (A)** Representative brightfield images of the organoids treated with DBZ, DAPT, IWP-2 or XAV939 at the end of the experiment (day 6). Scale bars, 500 uM, 200 uM, 100 uM from left to right. Black arrowheads point at folded organoids. **(B)** UMAP projections of scRNA-seq data from all organoid samples. **(C)** UMAP projections of scRNA-seq data coloured by days after hormonal stimulation, individual, hormonal stimulation, inhibitor used, phase of the cell cycle and sample ID. **(D)** qRT-PCR analysis for genes expressed in ciliated or secretory cells of hormonally-stimulated organoids treated with NOTCH inhibitors (blue) or WNT inhibitors (pink). Shown are the mean with SD levels of expression relative to housekeeping genes and control conditions (without inhibitors, black) at final time point of differentiation d6. n=3 patients. **(E)** Radial representation of the cosine distances similarity for single cells from organoid cultures to the centroids of *in vivo* epithelial cell types.

**Supplementary Tables:**

**Table S1.**

Metadata of samples, including biopsies and deceased transplant donors for scRNA-seq and visium analysis.

Single-cell and visium data:

| **Donor ID** | **Age (yo)** | **Cycle** | **Clinical information** | **10x kit** | **Cell cycle determination** | **Race** | **BMI** |
|---|---|---|---|---|---|---|---|
| A10 (discarded) | 35 | Proliferative | Recent possible miscarriage | 3’ v2 | | White | 24 |
| A13 | 37 | Proliferative | | 3’ v2 and visium | Histology | White | 18 |
| A16 (discarded) | 25 | Mid/late proliferative | | 3’ v2 | Histology | White | 24 |
| A30 | 23 | Early secretory | | 3’ v2 and visium | Histology | | 23 |
| E1 | 29 | Secretory (mid-late / day 25) | | 3’ v3 | LMP | | |
| E2 | 26 | Secretory (mid/ day 20) | Potential endometriosis | 3’ v3 | Histology + reported LMP | | |
| E3 | 26 | Proliferative (day 8) | Potential endometriosis | 3’ v3 | Histology + reported LMP | | |
| Trv2 | | Unknown | | 3’ v3 | | | |
| Trv3 | | Proliferative | | 3’ v3 | Histology | | |
| Trv4 | | Proliferative | | 3’ v3 | Histology | | |
| Trv5 | | Proliferative | | 3’ v3 | Histology | | |
| SAMN15049042 * | | Proliferative | | 3’ v3 | Computational and LMP | | |
| SAMN15049043 * | | Secretory (early) | | 3’ v3 | Computational and LMP | | |

| | | | | | |
|---|---|---|---|---|---|
| SAMN15049044 * | | Secretory (early) | | 3’ v3 | Computational and LMP |
| SAMN15049045 * | | Secretory (late) | | 3’ v3 | Computational and LMP |
| SAMN15049046 * | | Secretory (early-mid) | | 3’ v3 | Computational and LMP |
| SAMN15049047 * | | Secretory (early-mid) | | 3’ v3 | Computational and LMP |
| SAMN15049048 * | | Secretory (early-mid) | | 3’ v3 | Computational and LMP |
| SAMN15049049 * | | Secretory (mid) | | 3’ v3 | Computational and LMP |
| SAMN15049050 * | | Proliferative | | 3’ v3 | Computational and LMP |
| SAMN15049051 * | | Secretory (early) | | 3’ v3 | Computational and LMP |

* Data downloaded from (*21*)

In addition, four proliferative and four secretory endometrial samples were used for immunohistochemistry (IHC). These are from total hysterectomies from a tissue archive in the Centre for Trophoblast Research. All the samples chosen for IHC were histologically normal.

**Table S2. Table_S2.xlsx (separate file)**

Excel file containing quality control uterine atlas

**Table S3. Table_S3.xlsx (separate file)**

**(1)** Table containing differentially expressed genes in the main epithelial subpopulations detected in scRNAseq dataset. **(2)** Table containing Dorothea TF activities estimated in the epithelial subpopulations in scRNAseq dataset. Each row represents a TF/cluster analysis. **(3)** TFs predicted to be both differentially active and expressed.

**Table S4. Table_S4.xlsx (separate file)**

**(1)** Table containing differentially expressed genes in the main epithelial clusters detected in Visium dataset. **(2)** Table containing Dorothea results in Visium. Each row represents a TF/cluster analysis. **(3)** TFs predicted to be both differentially active and expressed.

**Table S5. Table_S5.xlsx (separate file)**

**(1)** Differentially expressed genes in the main and epithelial subsets in vivo. **(2-5)** Table containing CellPhoneDB interactions (epithelial-epithelial & fibroblasts epithelial) predicted for the lumenal, functional proliferative, functional secretory and basal microenvironments. Columns represent interaction cell pairs, rows represent the interactions. Table is binary, with 1 indicating that all the members of the interaction are expressed in at least 10% cells and at least one member is a DEGs in any cell-type in the pair with a positive log-fold change and FDR < 0.001.

**Table S6.**

Metadata of samples to derive endometrial organoids.

| **Donor ID** | **Age (yo)** | **Cycle** | **Clinical information** | **10x kit** |
|---|---|---|---|---|
| E001 | 32 | proliferative | ICSI for infertility of male partner | 3' v3 |
| B044 | 28 | secretory | IVF for subfertility of male partner | 3' v3 |
| B080 | N/A | N/A | N/A | 3' v3 |

**Table S7. Table_S7.xlsx (separate file).**

Quality control details of scRNA-seq dataset obtained from organoid cultures, which detail the list of samples obtained with hormone and inhibitor treatments.

**Table S8. Table_S8.xlsx (separate file).**

(1) Cosine distances defined by the transcriptome expression of single cells in organoid cultures compared to centroids defined by the epithelial cell type identified in the primary tissue. The log transformed expression from 1569 genes that were selected among the variable genes to be cell-type specific in cell types defined in the primary tissue. (2) Logistic regression predictions on the *in vivo* epithelial cells from the organoid populations without inhibitors that similarly used the expression from the same genes.

**Table S9. Table_9.xlsx (separate file).**

(1) Table containing differentially expressed genes in the organoid experiment without inhibitors detected by scRNAseq. (2) Table containing Dorothea TF activities estimated for the organoid subpopulations. (3-4) Transcription Factor comparison between epithelial subpopulations in vivo and organoid experiment without inhibitors for the glandular and ciliated lineages, respectively.

**Table S10. Table_S10.xlsx (separate file).**

(1) Table containing the cell counts and corresponding proportions identified as estrogen-induced, pre-ciliated, ciliated or secretory in the organoids on day 2 and 6, in relation to the 3 genotypes and inhibitor treatments considered targeting NOTCH and WNT. (2) Table containing the fold change for the frequency of a given cell type observed from the influence of an inhibitor treatment. The statistical significance of detected changes was reported for each 3 genotype independently using hypergeometric test, and robustness of detected effects across genotypes was then evaluated using a paired t-test for the 3 efficiencies reported without inhibitor treatment against the 3 reported with NOTCH and WNT inhibitors (2b and 2c respectively)

**Table S11. Table_S11.xlsx (separate file).**

(1) Cosine distances defined by the transcriptome expression of single cells in organoid cultures under inhibitor treatments compared to centroids defined by the epithelial cell type identified in the primary tissue. The log transformed expression from 1460 genes that were selected among the variable genes to be cell-type specific in cell types defined in the primary tissue. (2) Logistic regression predictions on the *in vivo* epithelial cells from the organoid populations with inhibitors that similarly used the expression from the same genes.

**Table S12. Table_S12.xlsx (separate file).**

(1) Table containing two pairwise differential expression analysis in the organoid experiment with inhibitors: i) measuring E+P effect in Secretory WNTi where *Secretory_WNTi* population is compared against *NH_WNTi*; and ii) measuring WNTi effect in Secretory lineage where *Secretory_WNTi* population is compared against *Secretory_Ctrl*. (2) Table containing Dorothea TF activities estimated from DEGs in (1).

**Table S13**

Antibodies used for immunohistochemistry

| **Primary antibodies** | | | | | | | |
|---|---|---|---|---|---|---|---|
| **Target** | **Clone** | **Product number** | **Supplier** | **Host species** | **Clonality** | **Dilution** | **Antigen retrieval buffer** |
| EPCAM | - | ab71916 | Abcam | Rabbit | Polyclonal | 1:1,500 | BOND ER2, pH 9.0 |
| Acetyl-alpha-Tubulin (Lys40) | 6-11B-1 | 12152S | Cell Signalling Technologies | Mouse | Monoclonal | 1:1000, 1:100 | AR pH 9.5 buffer |
| FOXJ1 | - | HPA005714 | Atlas | Rabbit | Polyclonal | 1:250 | AR pH 6.4 buffer |
| p73 | EP436Y | ab40658 | Abcam | Rabbit | Monoclonal | 1:750 | AR pH 9.5 buffer |
| HEY1 | - | ab22614 | Abcam | Rabbit | Polyclonal | 1:50 | AR pH 9.5 buffer |
| Glycodelin (PAEP) | EP870Y | ab 53289 | Abcam | Rabbit | Monoclonal | 1:500, 1:100 | AR pH 9.5 buffer |
| MMP7 | 111433 | MAB9071 | R&D Systems | Mouse | Monoclonal | 1:1000 | AR pH 6 buffer |
| ITGB6 | - | HPA023626 | Atlas Antibodies | Rabbit | Polyclonal | 1:450 | AR pH 6 buffer |
| PDPN | - | HPA007534 | Atlas Antibodies | Rabbit | Polyclonal | 1:200 | AR pH 6 buffer |

| **Secondary antibodies** | | | | | | |
|---|---|---|---|---|---|---|
| **Target** | **Conjugate** | **Product number** | **Supplier** | **Host species** | **Clonality** | **Dilution** |
| Rabbit IgG | HRP | G21234 | Thermo Fisher | Goat | Polyclonal | 1:1,500 |
| Rabbit IgG | Biotin | BA-1000 | Vector | Goat | Polyclonal | 1:200 |
| Mouse IgG | Biotin | BA-2000 | Vector | Horse | Polyclonal | 1:160 |

**Table S14**

Probes used for smFISH

| Target gene | Probe | Supplier | Product number |
|---|---|---|---|
| *NOTCH2* | RNAscope 2.5 LS Probe Hs-NOTCH2 | ACD, Bio-Techne | 488108 |
| *PAEP* | RNAscope 2.5 LS Probe Hs-PAEP | ACD, Bio-Techne | 453528 |
| *WNT7A* | RNAscope 2.5 LS Probe Hs-WNT7A-C2 | ACD, Bio-Techne | 408238-C2 |

**Table S15**

Taqman probes used for RT-qPCR

| Gene name | Description | Assay ID | Dye | Supplier | Product number |
|---|---|---|---|---|---|
| *FOXJ1* | Forkhead box J1; role in production of motile cilia | HS00230964_M1 | FAM | Thermo Fisher | 4331182 |
| *PIFO* | Primary cilia formation | HS00699083_M1 | FAM | Thermo Fisher | 4331182 |
| *PAEP* | Progestagen associated endometrial protein (glycodin) | Hs01046125_M1 | FAM | Thermo Fisher | 4331182 |
| *SPP1* | Secreted phosphoprotein 1 (osteopontin) | Hs00959010_M1 | FAM | Thermo Fisher | 4331182 |
| *HPRT1* (control) | Hypoxanthine phosphoribosyltran sferase 1 | Hs02800695_M1 | FAM | Thermo Fisher | 4331182 |
| *TBP* (control) | TATA-box binding protein | Hs00427620_M1 | FAM | Thermo Fisher | 4331182 |
| *TOP1* (control) | Topoisomerase (DNA) I | Hs00243257_M1 | FAM | Thermo Fisher | 4331182 |

**Table S16. Table_S16.xlsx (separate file).**

(1) New manually curated TF-target interactions. (2) Table containing TF regulons and scores used to quantify TF activities from DEGs.

**Table S17. Table_S17.xlsx (separate file).**

Table containing new curated cellphoneDB interactions.

**Table S18**

Components of ExM for culturing human endometrial organoids.

| **Product** | **Description** | **Company** | **Product Number** | **Final Concentration** |
|---|---|---|---|---|
| Advanced DMEM/F12 | Basal medium with reduced Fetal Bovine Serum (FBS) supplementation | Life Technologies | 12634010 | 1× |
| ALK-4, -5, -7 inhibitor, A83-01 | Inhibitor of TGF-beta type I receptor ALK5, ALK4, ALK7 kinases | System Biosciences | ZRD-A8-02 | 500 nM |
| B27 | Serum-free supplement minus vitamin A | Life Technologies | 12587010 | 1× |
| L-glutamine | Amino-acid | Life Technologies | 25030-024 | 2 mM |
| N2 | Serum-free supplement | Life Technologies | 17502048 | 1× |
| N-Acetyl-L-cysteine | Antioxidant and mucolytic agent | Sigma-Aldrich | A9165-5G | 1.25 mM |
| Nicotinamide | Supplement amide derivative of vitamin B3 and a PARP inhibitor | Sigma-Aldrich | N0636 | 10 nM |
| Primocin | Antimicrobial agent | Invivogen | ant-pm-1 | 100 µg/ml |
| Recombinant human EGF | Epidermal Growth Factor | Peprotech | AF-100-15 | 50 ng/ml |

| | | | | |
|---|---|---|---|---|
| Recombinant human FGF-10 | Fibroblast Growth Factor-10 | Peprotech | 100-26 | 100 ng/ml |
| Recombinant human HGF | Hepatocyte Growth Factor | Peprotech | 100-39 | 50 ng/ml |
| Recombinant human Noggin | Bone Morphogenetic Protein 4 antagonist | Peprotech | 120-10c | 100 ng/ml |
| Recombinant human Rspondin-1 | Wnt/beta-catenin signaling pathway stimulator | Peprotech | 120-38 | 500 ng/ml |
| Components added to single cells. | | | | |
| A 83-01 | Inhibitor of Transforming Growth Factor beta kinase type 1 | Sigma-Aldrich | SML0788 | 1 µg/ml |
| CHIR 99021 | Inhibitor of Glycogen Synthase Kinase 3 (GSK-3) | Tocris | 4423 | 0.2 µg/ml |
| Hormones used for differentiation of human endometrial organoids | | | | |
| 8-Bromoadenosine 3’5’-cyclic monophosphate (cAMP) | Protein kinase A activator | Sigma-Aldrich | B7880 | 1 µM |
| Progesterone (P4) | Steroid hormone produced by the corpus luteum | Sigma-Aldrich | P7556 | 1 µM |
| Prolactin (PRL) | Neuroendocrine hormone secreted by the pituitary gland | Peprotech | 100-07 | 20 ng/ml |
| B-estradiol (E2) | Steroid hormone produced by the corpus luteum | Sigma-Aldrich | E4389 | 10 nM |

| | | | | |
|---|---|---|---|---|
| Inhibitors used for signalling inhibition | | | | |
| DBZ | Gamma-secretase inhibitor; inhibits Notch pathway | Tocris | 4489 | 1 µM |
| DAPT | Gamma-secretase inhibitor; inhibits Notch pathway | Tocris | 2634 | 20 µM |
| XAV939 | Tankyrase inhibitor; inhibits WNT pathway | Tocris | 3748 | 2 µM |
| IWP-2 | PORCN inhibitor; inhibits Wnt processing and secretion | Tocris | 3533 | 2 µM |
